# High-dimensional mediation analysis to elucidate the role of metabolites in the association between PFAS exposure and reduced SARS-CoV-2 IgG in pregnancy

**DOI:** 10.1101/2024.12.16.628663

**Authors:** Haibin Guan, Jia Chen, Kirtan Kaur, Bushra Amreen, Corina Lesseur, Georgia Dolios, Syam S. Andra, Srinivasan Narasimhan, Divya Pulivarthi, Vishal Midya, Lotje D. De Witte, Veerle Bergink, Anna-Sophie Rommel, Lauren M. Petrick

## Abstract

We previously found that per- and polyfluoroalkyl substances (PFAS) mixture exposure is inversely associated with SARS-CoV-2 IgG (IgG) antibody levels in pregnant individuals. Here, we aim to identify metabolites mediating this relationship to elucidate the underlying biological pathways. This cross-sectional study included 59 pregnant participants from a US-based pregnancy cohort. Untargeted metabolomic profiling was performed using Liquid Chromatography-High Resolution Sass spectrometry (LC-HRMS), and weighted Quantile Sum (WQS) regression was applied to assess the PFAS and metabolites mixture effects on IgG. Metabolite indices positively or negatively associated with IgG levels were constructed separately and their mediation effects were examined independently and jointly. The PFAS- index was negatively associated with IgG levels (beta=-0.315, p<0.001), with PFHpS and PFHxS as major contributors. Two metabolite-indices were constructed, one positively (beta=1.249, p<0.001) and one negatively (beta=-1.200, p<0.001) associated with IgG. Key contributors for these indices included trigonelline, adipate, p-octopamine, and n-acetylproline. Analysis of a single mediator showed that 74.6% (95% CI: 45.9%, 98.0%) and 68.6% (95% CI: 41.8%, 94.1%) of the PFAS index-IgG total effect were mediated by the negative and positive metabolites-indices, respectively. Joint analysis of the metabolite-indices indicated a cumulative mediation effect of 83.8% (95% CI: 58.1%, 98.7%). Enriched pathways associated with these metabolites indices were phenylalanine, tyrosine, and tryptophan biosynthesis and arginine metabolism. We observed significant mediation effects of plasma metabolites on the PFAS-IgG relationship, suggesting that PFAS is associated with alteration in the balance of plasma metabolites that contributes to reduced plasma IgG production.

## INTRODUCTION

PFAS (Per- and Polyfluoroalkyl Substances), often called “forever chemicals,” are industrial chemicals widely used due to their durability. Their extensive use and tendency to bioaccumulate have raised significant public health concerns. In particular, PFAS exposure has been associated with weakened immune responses after vaccination, including reduced immunosuppression biomarkers such as immunoglobulin production (Luoping Zhang. et al., 2022). Studies have highlighted the connection between PFAS exposure during pregnancy and immune response in both mothers and children. For example, the Norwegian Mother and Child Cohort Study reported significant inverse associations between the concentrations of maternal blood perfluorooctanoic acid (PFOA), perfluorooctanesulfonic acid (PFOS), perfluorohexanesulfonic acid (PFHxS), and perfluorononanoic acid (PFNA) measured at delivery, and the levels of anti-rubella antibodies in the children’s sera at age three years (Granum B et al., 2013). Given PFAS’s known influence on immune function, recent studies have begun exploring how PFAS exposure may impact immune responses to specific infections, including SARS-CoV-2. Indeed, communities exposed to higher levels of PFAS exhibited higher odds of COVID-19 mortality (Catelan D et al., 2021). Moreover, elevated plasma levels of perfluorobutanoic acid (PFBA) are associated with more severe SARS-CoV-2 infections (P Grandjean, 2020), while higher serum levels of PFOS, PFHxS, and PFNA are linked to a weaker antibody response after SARS-CoV-2 infection (Hollister J, 2023). Our team previously found that levels of a mixture of PFAS, including linear- (n-)PFOA, PFHxS, perfluoroheptanesulfonic acid (PFHpS), and perfluorohexanoic acid (PFHxA), were inversely associated with maternal SARS-CoV-2 IgG (IgG) antibody levels in pregnant individuals (K. Kaur et al., 2023). However, the underlying biological mechanisms are not yet fully understood.

Metabolomics, representing the final products of biochemical changes and closest omics layer to disease and health phenotypes, has advanced rapidly with developments of high-resolution mass spectrometry (HRMS) which can simultaneously measure tens of thousands of endogenous and exogenous metabolites representing a range of biologically relevant pathways for discovery. To understand the disease’s pathophysiology and potential therapeutic strategies, several studies have investigated the immune response and metabolic signatures in SARS-CoV-2 patients (Song JW, et al., 2020; Thomas T, et al., 2020; Gardinassi LG,et al., 2020; Takeshita H, et al., 2022; Valdés A, et al.,2022). Furthermore, as endogenous metabolites have previously been associated with PFAS exposures (India-Aldana S, et al., 2023), disrupted biological pathways are considered as potential mediators linking exposures to diverse health outcome (Wei, M., et al., 2021; Mallol R, et al., 2021).

Traditional mediation analysis methods (Baron and Kenny, 1986) are designed to decompose the total effect of an exposure on an outcome into direct and indirect effects through mediators. However, these methods are generally limited to single-exposure, single-mediator scenarios (Valeri and VanderWeele, 2013; Daniel RM, et al., 2015) and struggle with high-dimensional data, particularly when mediators are highly correlated. In studies involving numerous, interrelated metabolites, individual mediation analysis is not only computationally expensive but also risks inflating the multiple comparison error. This complexity underscores the need for an approach that can efficiently summarize multiple correlated metabolites into an “interpretable index,” capturing their joint effects on the outcome. For instance, several comprehensive approaches have been developed to focus on generating latent factors through dimensionality reduction techniques, such as principal component analysis (PCA) (Huang and Pan, 2016; Goodrich, et al., 2024) and partial least squares (PLS) (Assi N, et al., 2018). These methods transform correlated mediators into independent latent factors that may represent molecular pathways, followed by mediation analysis on these factors. Weighted Quantile Sum (WQS) regression (Carrico, C, et al., 2013) as a mixture modeling tool offers a practical and effective solution by aggregating metabolites into indices, thus reducing dimensionality while preserving interpretable insights. WQS can efficiently handle both positive and negative metabolites- indices, making it particularly suited to our study, where metabolite levels can be increasing or decreasing in the link between PFAS exposures and IgG levels. By creating these mixture indices, WQS not only reduces the high computational burden but also addresses the challenge of multiple comparisons, allowing for a robust assessment of the overall metabolite effects within a comprehensive framework.

In this study, we profiled the plasma metabolome in the same individuals as our previous study (K. Kaur et al., 2023) and conducted mediation analysis. We applied WQS in a mediation analysis integrating high-dimensional plasma metabolomics and PFAS mixture exposures to provide insights on the biological impacts of PFAS exposures linked to IgG response in pregnant individuals.

## MATERIALS AND METHODS

### Study Population

The Generation C cohort was created to explore the impact of SARS-CoV-2 infection on pregnancy and birth outcomes in pregnant individuals. A total of 3,157 pregnant individuals were enrolled from April 20, 2020 until Feb 24, 2022. They met the eligibility criteria of being 18 years or older, receiving obstetrical care at Mount Sinai Health System, having a single live birth, and residing in NYC. As part of their routine obstetrical care, blood samples were collected from participants, following informed consent. Past SARS-CoV-2 infection was verified through a serological IgG antibody test and a review of their electronic medical records (EMR) (Gigase et al., 2024; Janevic et al., 2022; Lesseur et al., 2022). Procedures were approved by the institutional review board at the Icahn School of Medicine at Mount Sinai. We previously reported an inverse relationship between PFAS and IgG levels in 72 participants with available PFAS and IgG antibody data (K. Kaur et al., 2023). For the follow-up study, we measured the metabolome in a subset of these individuals with sufficient plasma remaining (N=59).

### Maternal Plasma PFAS Measurements

Maternal plasma PFAS measurements have been described previously (K. Kaur et al., 2023). In brief, maternal plasma samples, were stored at −80°C and analyzed for 16 PFAS congeners using a low-volume (100 μL) and high-sensitivity (0.2 ng/mL LOD) isotope-dilution LC-MS/MS assay at the Icahn School of Medicine at Mount Sinai’s Human Health Exposure Analysis Resource (HHEAR) Targeted Analysis Laboratory. The laboratory adheres to internal quality assurance protocols and participates in proficiency testing programs such as G-EQUAS (https://app.g-equas.de/web/) (Göen T et al., 2012) and CTQ-AMAP (https://www.inspq.qc.ca/en/ctq/eqas/amap/description) (CTQ, AMAP: AMAP Ring Test for Persistent Organic Pollutants in CTQ, 2022). The method, based on CDC protocols with modifications, quantifies widely studied PFAS (n-PFOS, branched- (Sm-)PFOS, n-PFOA, PFHxS, PFNA, and perfluorodecanoic acid [PFDA]), emerging PFAS (perfluorobutanesulfonic acid [PFBS], PFHpS, perfluoroheptanoic acid [PFHpA], PFHxA, perfluoroundecanoic acid [PFUnDA], and perfluorododecanoic acid [PFDoDA]), and replacements (n-ethylperfluoro-1- octanesulfonamido acetic acid [N-EtFOSAA] and n-methylperfluorooctanesulfonamidoacetic acid (mixed isomers) [N-MeFOSAA]). For PFAS concentrations below the limit of detection (LOD), values were estimated as LOD/(√2). PFAS compounds detected below the LOD in more than 40% of samples were excluded. The final analysis focused on nine PFAS compounds: PFOS (n- and Sm- PFOS combined), PFHxA, PFHxS, n-PFOA, PFHpS, PFDA, PFNA, PFBS, and PFUnDA.

### SARS-CoV-2 IgG Antibody Quantification

Serological testing for quantifying IgG antibodies against the SARS-CoV-2 spike (S) protein was performed with an enzyme-linked immunosorbent assay (ELISA) developed at the Icahn School of Medicine at Mount Sinai (Stadlbauer et al., 2020). Results are reported as antibody units per milliliter (AU/mL).

### Maternal Plasma Metabolites Analysis

Plasma metabolomics was performed with LC-HRMS using established protocols (Hua X et al., 2023; Niedzwiecki MM et al., 2021; Yu M et al., 2024). For each sample, 50 µL of plasma was combined with 150 µL of ice-cold methanol crash solution, vortexed for 10 seconds, incubated at -80 °C for 30 minutes, and centrifuged at 13,000g for 15 minutes at 4 °C. The samples were then evaporated using a Savant SC250EXP SpeedVac vacuum concentrator at 35 °C for 90 minutes. Dried sample extracts were stored at -80 °C. Pooled quality control (QC) samples were created by combining 20 μL extracts from each plasma sample and re-aliquoting. In the same extraction process, solvent blanks were prepared by substituting plasma with water. The samples were analyzed in a randomized sequence, with pooled QCs injected every 10 samples throughout the run. Procedural blank extracts were analyzed three times at the start and end of the batch. The samples were analyzed in one batch by ultra-performance liquid chromatography coupled to a HRMS in both reverse-phase positive (RPP) ionization and reverse-phase negative (RPN) ionization modes and Zwitterion Hydrophilic Interaction Chromatography (HILIC) chromatography in positive ionization mode (ZHP). Metabolite identifications were made using our local database, which comprises over 1000 biologically and environmentally relevant reference standards analyzed under the same conditions. Matching involved considering retention time (± 8 sec), accurate mass (< 20ppm), isotope distribution, and MS/MS fragmentation pattern (when available) against the in-house Personal Chemical Database Library and Profinder software (Agilent Technologies, Santa Clara, USA), resulting in annotation confidence levels 1 and 2 (Schymanski EL, 2014). Metabolite classes were determined using ClassyFire (Djoumbou Feunang et al., 2016).

Annotated metabolites in each of the three modes were filtered and normalized. A 20% missing value threshold was set and metabolites exceeding this threshold across samples were excluded from further analysis. Only metabolites above a signal-to-noise threshold of 1.5 median signal fold change in samples compared to solvent blank were maintained. To correct the batch effects, all metabolites were corrected using TIGERr (Siyu Han et al., 2022). A quality criteria threshold of 30% coefficient of variation (CV) in repeated injections of pooled QC samples was set to eliminate metabolites with high variability. We then combined the cleaned and normalized metabolites of the three modes and eliminated any duplicates (see **Figure S1**), giving priority to the analysis mode with higher abundance and lower missingness. Subsequently, a log2 transformation was applied to the remaining metabolite values to normalize the data and facilitate statistical comparisons. We then imputed the missing values by using a 2- step mechanism with the package “Mechanism-Aware Imputation (MAI)” (Dekermanjian, J.P. et al, 2022). Finally, all values were standardized in z-score for downstream statistical analysis.

### Covariates

For each participant, demographic variables including maternal age, race/ethnicity, pre-pregnancy body mass index (BMI), gestational age at blood draw, parity, as well as the number of administered COVID-19 vaccine doses prior to blood draw. We also included the type of insurance as a measure of socioeconomic status. Cohort demographics and clinical characteristics of the 59 Generation C participants in this study are summarized in **Table 1**.

**Table 1.**
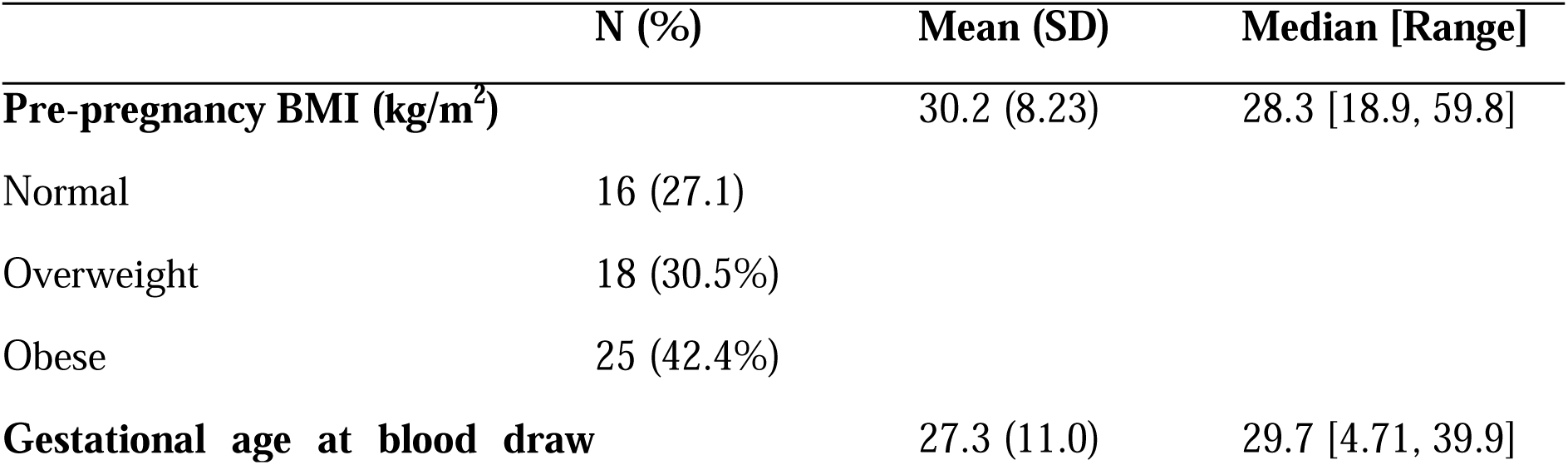
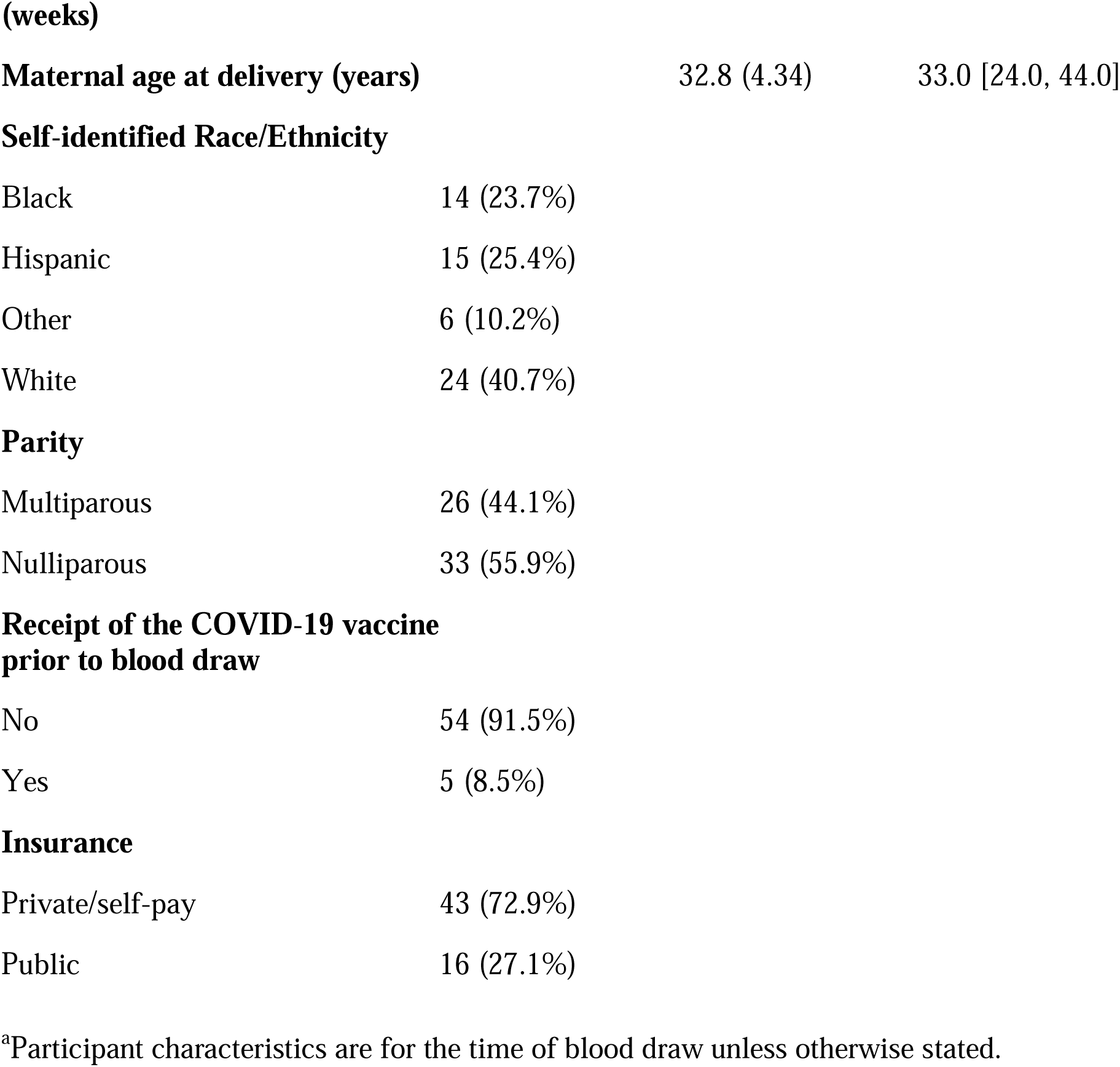
Descriptive characteristics of the population’s potential confounding variables in the current study (N=59)^a^.

### Statistical Analysis

In this study, all analyses were adjusted for maternal age, maternal race/ethnicity, parity, type of insurance, gestational age at blood draw (weeks), receipt of the COVID-19 vaccine before blood collection, and pre-pregnancy BMI. We first conducted univariate linear regression analysis for each individual PFAS congener to assess its association with IgG levels. Following this, we used WQS regression (Carrico, C, et al., 2013) to estimate the combined effect of the 9 PFAS exposure mixture on IgG levels in the adverse direction by creating a single outcome-related index, called the PFAS-index. This analysis was restricted to the negative direction based on our previous findings in a larger population of 72 individuals (K. Kaur et al., 2023). We then used a random-subset implementation of WQS (WQS_RS_) appropriate for highly collinear omics contexts (Curtin, P. et al., 2019) to identify an index and assess the combined impact of metabolites. Two indices were constructed to capture distinct associations between metabolite levels and IgG. The first index [metabolites-index (Neg)] includes metabolites whose increased plasma levels are associated with lower IgG, while the second [metabolites-index (Pos)] includes those whose increased levels are associated with higher IgG. This separation allows us to capture both positive and negative relationships, reflecting potentially distinct biological pathways. In this step, each metabolite within the index was weighted based on its contribution to the observed association, which was estimated through bootstrapping (n=3,000).

To further examine whether either direction of the metabolites-index could potentially mediate the association between the PFAS-index and the IgG antibody levels, we implemented the mediation analysis using the R package “lavaan”. In the single mediator analysis, we regressed the metabolites-index on the PFAS-index and covariates in the mediator model, and regressed IgG antibody levels on the PFAS-index, metabolites-index, and covariates in the outcome model. Together, these models estimate the direct and indirect effects of the PFAS-index on IgG levels. To study the combined effect of the two metabolic-indices, we also employed a multiple-mediator analysis. Rather than separately estimating the weights in the WQS step and fixing them in the mediation model, we integrated the WQS and mediation analyses within a single iterative bootstrap procedure with 5000 simulations to account for uncertainty in weight estimation. To obtain robust standard errors for the mediation estimates, we employed maximum likelihood estimation with Huber-White robust standard errors (Huber, 1981; White, 1980). **Figure S2** is a schematic of the mediation analysis workflow conducted in this study, detailing both univariate and multivariate models that assess the direct and indirect effects of PFAS exposure on IgG levels through specific metabolite indices.

To further interpret the findings functionally and identify pathways potentially involved in the mediation process, we performed metabolite set enrichment analyses. We focused on the set of metabolites that passed the weight threshold (1/metabolite set size of 286) in both metabolites- indices, which reflects the key metabolites contributing to the observed associations with IgG levels. Over Representation Analysis (ORA) was performed using MetaboAnalystR 4.0 (Pang, Z., 2024) with the human Kyoto Encyclopedia of Genes and Genomes (KEGG) pathway database. The Enrichment Ratio, reported in our results, represents the proportion of significant metabolites within a given pathway relative to the expected proportion under the null hypothesis. The significance of the enriched pathway was determined using Fisher’s exact test, with statistical significance set at a p-value threshold of 0.05.

## RESULTS

Demographic and clinical characteristics of the study population (N=59) are found in Table 1. The demographic characteristics of this population (N=59) are similar to those in our previous study (N=72). At the time of the blood collection, the average age of the mother was 32.8 years (SD=4.34) and the average gestational age at blood draw was 27.3 weeks (SD=11.0). The participants were nulliparous (55.9%) and multiparous (44.1%). The population included 23.7% (n=14) self-identified as Black, 25.4% (n=15) as Hispanic, 40.7% (n=24) as white, and 10.2% (n=6) as other. Most participants were privately insured (72.9%). Five participants had received at least one COVID-19 vaccine dose before blood collection.

Following data preprocessing, 286 annotated metabolites were selected for downstream analysis. Summary statistics for those 286 metabolites, based on their log₂-transformed abundances and grouped by metabolite superclass, are presented in **Tables S1** and **Table S2**. Metabolites demonstrated high collinearity (**Figure S3**). Additionally, descriptive statistics for the PFAS-index and two metabolite indices are reported in **Table S3**. The results of linear regression analysis for individual PFASs are provided in the supplementary material (see **Figure S4**).

In the current study, the PFAS-index was inversely associated with IgG (Total Effect= -0.315 [95% CI: -0.478, -0.175]), see **Supplementary Table S4**, **Figure S4**). Specifically, a one-quartile increase in the PFAS mixture was associated with a decrease in the IgG antibody level by 10^0.315^ AU/mL.

In the analysis of a single mediator with metabolites-index (Neg), we found a significant indirect effect. Specifically, for each one-quartile increase in the PFAS-index, the metabolites- index (Neg) increased by 0.223 units (beta=0.223 [95% CI: 0.108, 0.347], p<0.001), resulting in a subsequent decrease in the IgG antibody level by 10^1.122^ AU/mL (beta=-1.122 [95% CI: - 1.490, -0.797], p<0.001) (see **Figure 1.a-c**). As a result, 74.6% (95% CI: 45.9%, 98.0%) of the total effect of the PFAS-index on decreased IgG levels could be explained by the indirect effect of the metabolites-index (Neg) (Indirect Effect=-0.248 [95% CI: -0.404, -0.1117], p<0.001).

**Figure 1.**
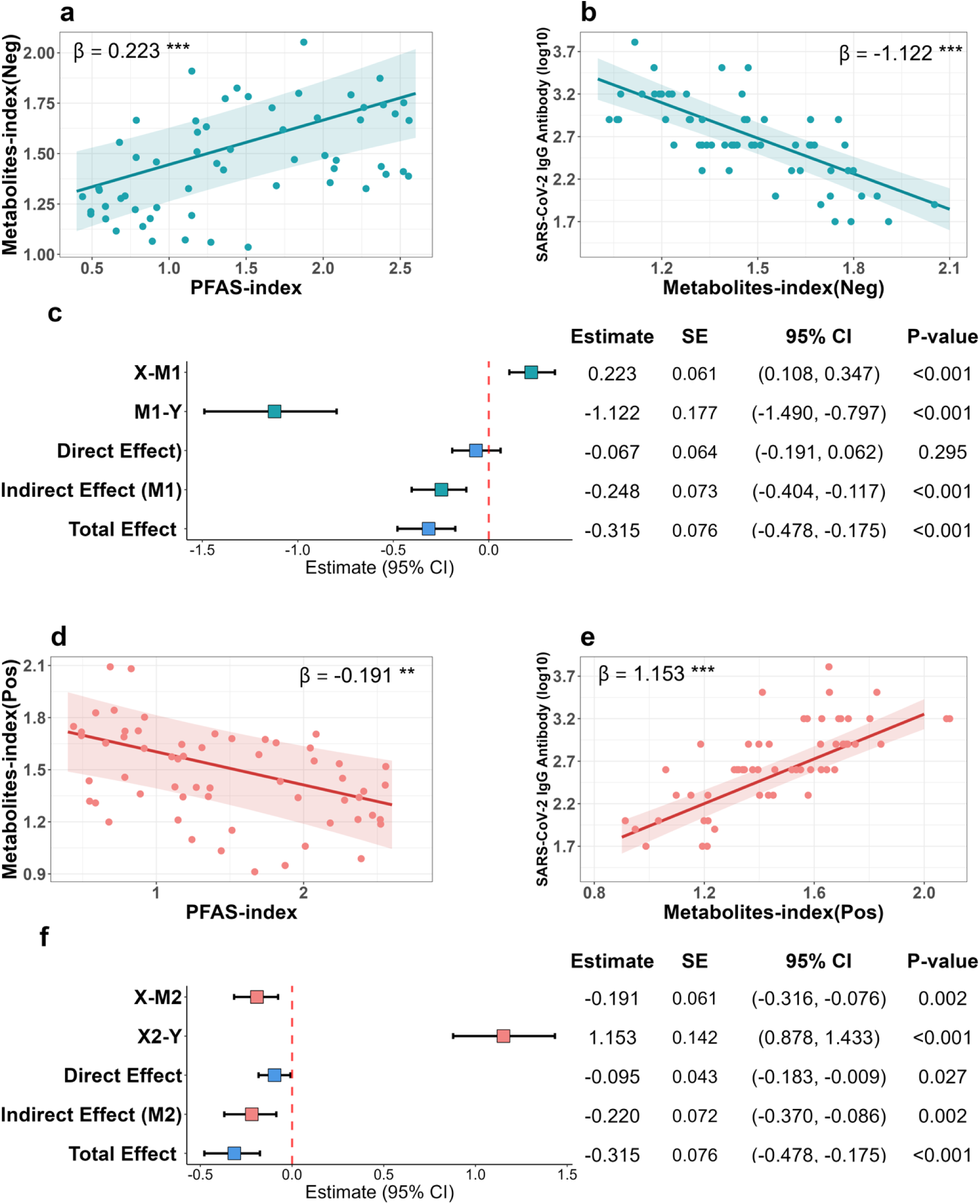
Results of single mediator analyses for PFAS-index, metabolites-index (Neg) or metabolites-index (Pos), and SARS-CoV-2 IgG antibody levels (N = 59) using models adjusted for maternal age, maternal race/ethnicity, parity, type of insurance, gestational age at blood draw (weeks), receipt of the COVID-19 vaccine before blood collection, and pre-pregnancy BMI. 95% confidence levels (CIs) are derived from the 5,000 bootstrap simulations. **a)** Scatter plot of the estimated beta (adjusted for the model residual when covariates are included in the model) for the association between the metabolites-index (Neg) and PFAS-index from the mediator model. **b)** Scatter plot of the estimated beta (adjusted for the model residual when covariates are included in the model) for the association between IgG levels and metabolites-index (Neg) from the outcome model. **c)** Forest plot of PFAS-index, metabolites- index (Neg) and SARS-CoV-2 IgG antibody levels mediation analysis summary parameters. **d)** Scatter plot of the estimate estimated beta (adjusted for the model residual when covariates are included in the model) for the association between metabolites-index (Pos) and PFAS-index from the mediator model. **e)** Scatter plot of the estimated beta (adjusted for the model residual when covariates are included in the model) for the association between IgG levels and the metabolites-index (Pos) from the outcome model. **f)** Forest plot of PFAS-index, metabolites- index (Pos) and SARS-CoV-2 IgG antibody levels mediation analysis summary parameters. All p-values are estimated using a two-sided test for t-statistics from linear regression models. Statistical significance levels for the effects: * (p < 0.05); **(p < 0.01); and ***(p < 0.001). X (PFAS-index exposure); M1 (Metabolites-index [Neg]); M2 (Metabolites-index [Pos]); Y (SARS-CoV-2 IgG levels).

Likewise, in the analysis of a single mediator with metabolites-index (Pos), we found a significant indirect effect, showing that for each one-quartile increase in the PFAS-index, the metabolites-index (Pos) decreased by 0.191 units (beta=-0.191 [95% CI:-0.316, -0.076], p=0.002), resulting in a subsequent decrease in the IgG levels by 10^1.1s3^ AU/mL (beta=1.153 [95% CI: 0.878, 1.433], p<0.001) (see **Figure 1.d-f**). As a result, 68.6% (95% CI: 41.8%, 94.1%) of the total effect of the PFAS-index on IgG could be explained by the indirect effect of the metabolites-index (Pos) (Indirect Effect =-0.220 [95% CI: -0.370,-0.086], p=0.002).

In the joint mediation model, where both positive and negative metabolites-indices were included to control for the mutual influence, we found a significant but smaller indirect effect for both metabolites-indices, 36.9% (95% CI: 18.1%, 66.1%) for the metabolites-index (Neg) (Indirect Effect=-0.120 [95% CI: -0.224, -0.048], p=0.007) and 49.1% (95% CI: 27.7%, 72.1%) for the metabolites-index (Pos) (Indirect Effect=-0.157 [95% CI: -0.276, -0.059], p=0.005), resulting in a cumulative mediation effect of 83.8% (95% CI: 58.1%, 98.7%) (see **Figure 2a**).

**Figure 2.**
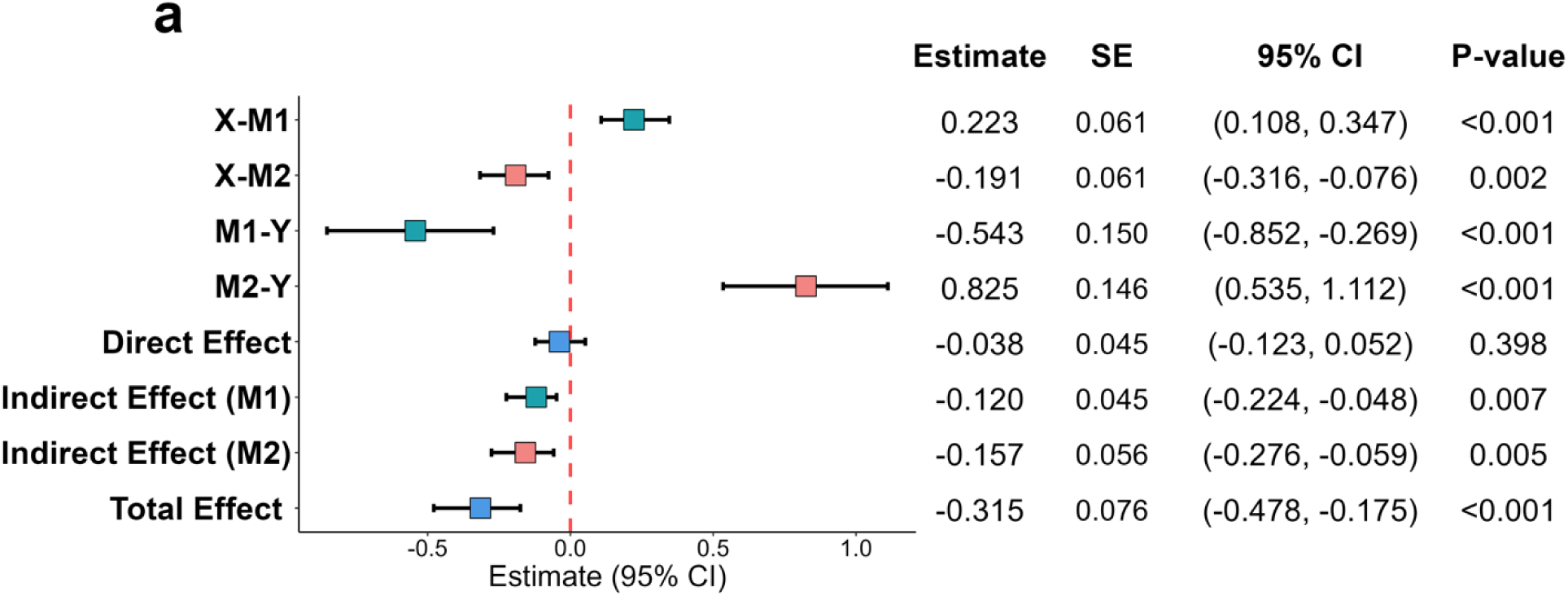
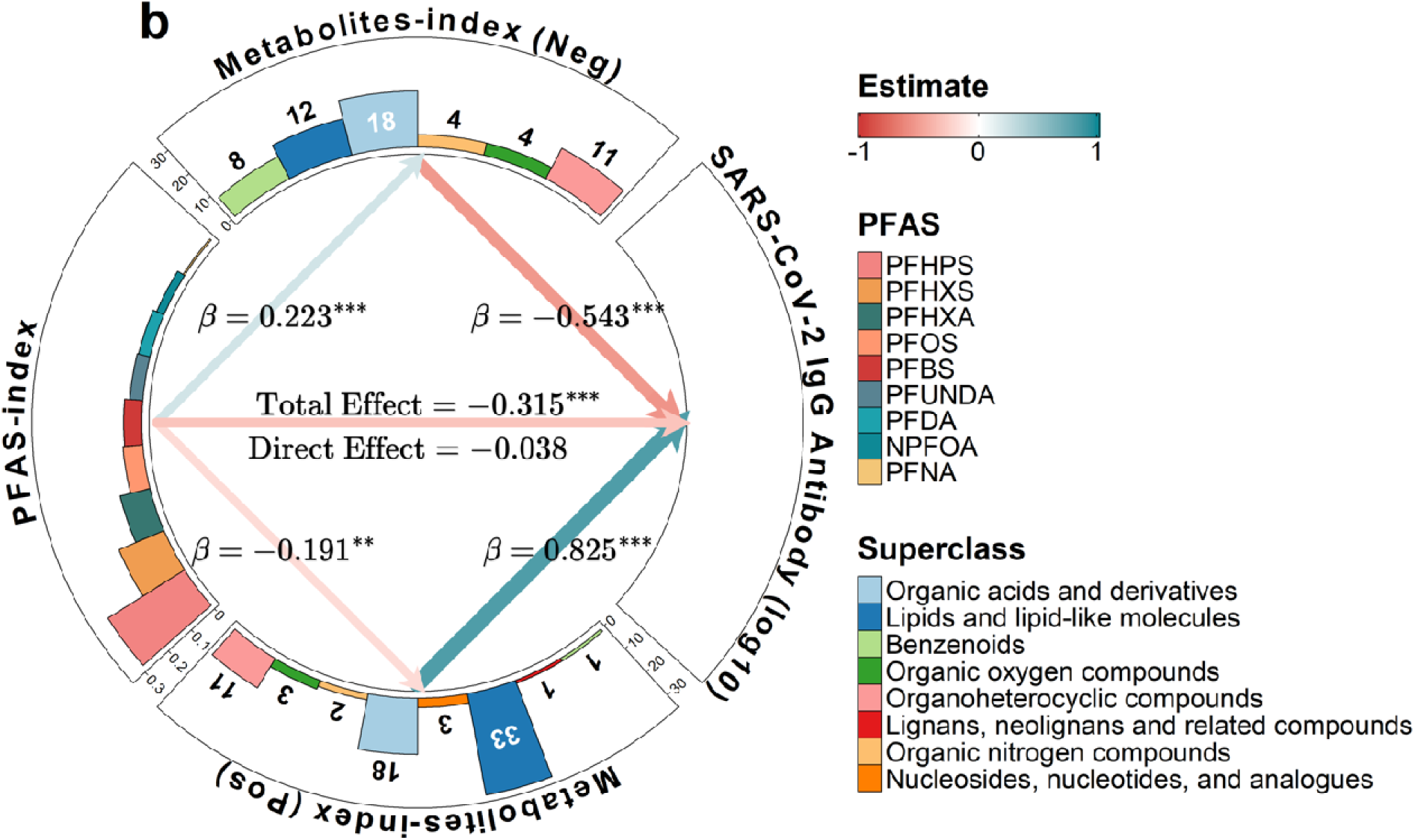
**a)** Forest plot of joint mediation analysis with two mediators, metabolites-index (Pos) and metabolites-index (Neg), on PFAS-index and SARS-CoV-2 IgG antibody levels (N = 59) adjusted for maternal age, maternal race/ethnicity, parity, type of insurance, gestational age at blood draw (weeks), receipt of the COVID-19 vaccine before blood collection, and pre-pregnancy BMI. 95% CIs are derived from the 5,000 bootstrap simulations. X (PFAS-index exposure); M1 (Metabolites-index [Neg]); M2 (Metabolites-index [Pos]); Y (SARS-CoV-2 IgG levels). **b)** Circos plot showing the estimates from the joint mediation analysis with two metabolites-indices on the PFAS-index and IgG levels. The thickness of the links between variables represents the strength of the estimates, with the color gradient ranging from red (-1) to turquoise (1) to depict the direction and magnitude of these relationships. Within the PFAS-index sector, the bar plot displays the estimated weights assigned to each PFAS component derived from the WQS models based on 5,000 bootstrap simulations, while the bar plots within the two metabolites-indices illustrate the number of chemical superclasses that exceeded the cutoff threshold within each metabolites-index. All p-values were estimated using a two-sided test for t-statistics from linear regression models, confidence interval. Statistical significance levels for the effects: * (p < 0.05); **(p < 0.01); and ***(p < 0.001).

In the WQS regression models adjusting for covariates and constrained to either the negative or positive direction, the metabolites-index (Pos) showed a significant association with IgG levels (mean beta = 1.249, SE = 0.140, p < 0.001). Specifically, a one-quartile increase in the metabolite mixture was associated with an increase in the IgG antibody level by 10^1.249^ AU/mL. In contrast, the metabolites-index (Neg) showed a significant association with IgG levels (mean beta = -1.200, SE = 0.160, p < 0.001) (see Supplementary **Table S4**), where a one-quartile increase in the metabolite mixture was associated with a decrease in the IgG antibody level by 10^-1.200^ AU/mL.

Figure S6 depicts the complete list of 57 and 72 metabolites that passed the cutoff threshold (1/286) in metabolites-indices (Neg) and (Pos), respectively. For clarity, only the estimated weights for the top 10 metabolites that passed the cutoff threshold (1/286) are depicted in Figure 3. Weights indicate the relative contribution of each metabolite to the index. Among the 286 metabolites, plasma p-octopamine is the most influential compound in the negative association with plasma IgG levels (median weight = 0.021) while plasma trigonelline is the primary driver in the positive association with plasma IgG levels (median weight = 0.016) (see Figure 3). Over half of the metabolites that passed the cutoff threshold in the metabolites-index (Neg) were classified as either organic acids and derivatives (18/57) or lipids and lipid-like molecules (12/57). Similarly, 45.8% of the metabolites surpassing the cutoff in the metabolites-index (Pos) were lipids and lipid-like molecules (33/72). Notably, 11 metabolites classified as organoheterocyclic compounds had relatively higher weights in this index, with 6 out of the 11 organoheterocyclic compounds ranking among the top 10 most weighted metabolites (see Figure 2b**, Figure S6**). Figure 4 showcases a comprehensive network diagram that further elucidates the relationships between key PFAS compounds and metabolites, along with their relative contributions to the PFAS-index and the two metabolites-indices.

**Figure 3.**
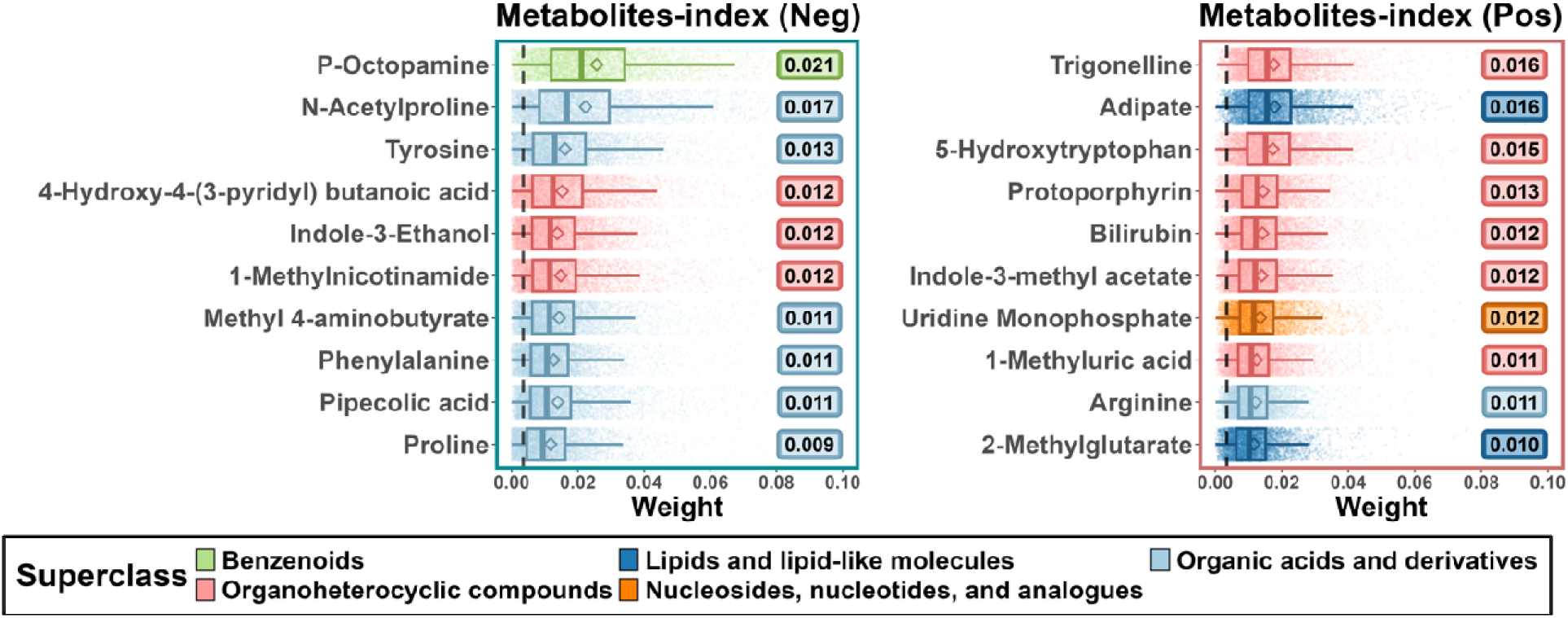
The box plot shows the weights of top 10 metabolites whose median weights exceeded the cutoff threshold in the metabolites-index (Neg) (left) and metabolites-index (Pos) (right), respectively. Metabolites are ranked based on their median weight, which is shown as a text label. The box-plot depict the 25^th^, 50^th^, and 75^th^ percentiles, while the whiskers extend to the 10^th^ and 90^th^ percentiles. Diamonds indicate the mean weights, and individual data points represent the estimated weights associated with the metabolites-indices (Neg) and (Pos) through the 5000 simulations. The dashed dark line represents the cutoff (1/286), the colors represent the respective chemical superclass of each metabolite.

**Figure 4.**
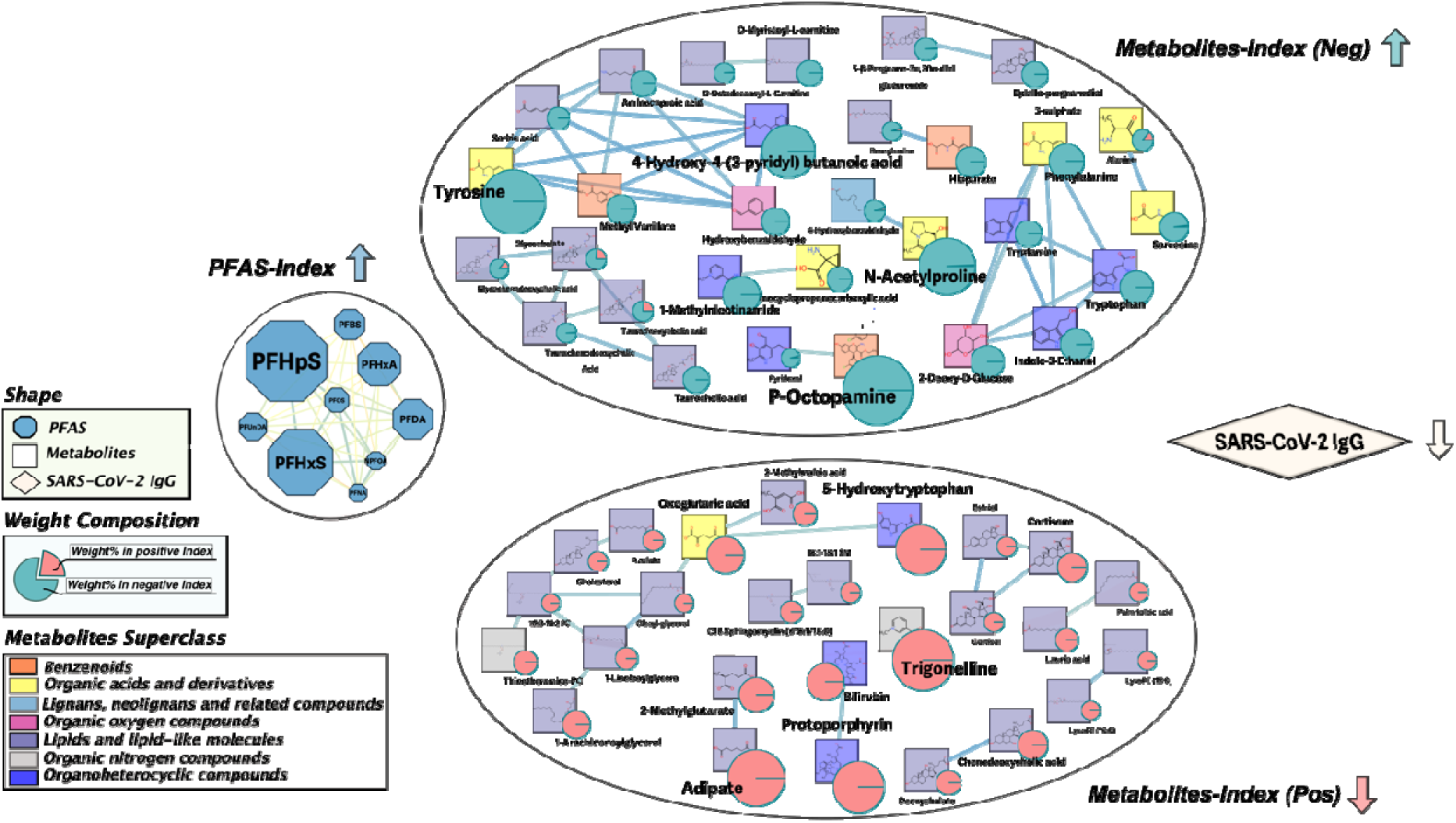
The network illustrates the primary contributors within the PFAS-Index and two metabolites-indices. Nodes represent different entities: metabolites (rectangles), PFAS exposure (octagons), and SARS-CoV-2 IgG levels (diamond). Metabolites are color-coded according to their chemical superclass, with pie chart size reflecting both their relative weight and composition across the indices. Links between metabolites and PFAS nodes represent their partial correlations.

To characterize the metabolic pathways associated with metabolites-index (Neg) and (Pos), we conducted ORA using the KEGG library. For the 57 metabolites in the metabolites-index (Neg), seven pathways were significantly enriched (Fisher’s p < 0.05) with phenylalanine, tyrosine, and tryptophan biosynthesis the most significant pathway (Figure 5). For the 72 metabolites in th metabolites-index (Pos), nine pathways were significantly enriched with arginine biosynthesis the most significant pathway (Figure 5). Given that ORA can be biased by the metabolite list used for analysis (i.e., 286 metabolites), we also considered the full set of identified pathways for discovery (**Figure S7**). Overall, the 57 metabolites that contributed to the metabolites-index (Neg) and the 72 metabolites that contributed to the metabolites-index (Pos) were associated with a total of 42 metabolic pathways. Among these, 16 pathways were shared between the two indices (e.g. primary bile acid biosynthesis, tryptophan metabolism and taurine and hypotaurine metabolism), while 8 and 18 pathways were unique to the metabolites-index (Neg) and metabolites-index (Pos), respectively (**Figure S7**).

**Figure 5.**
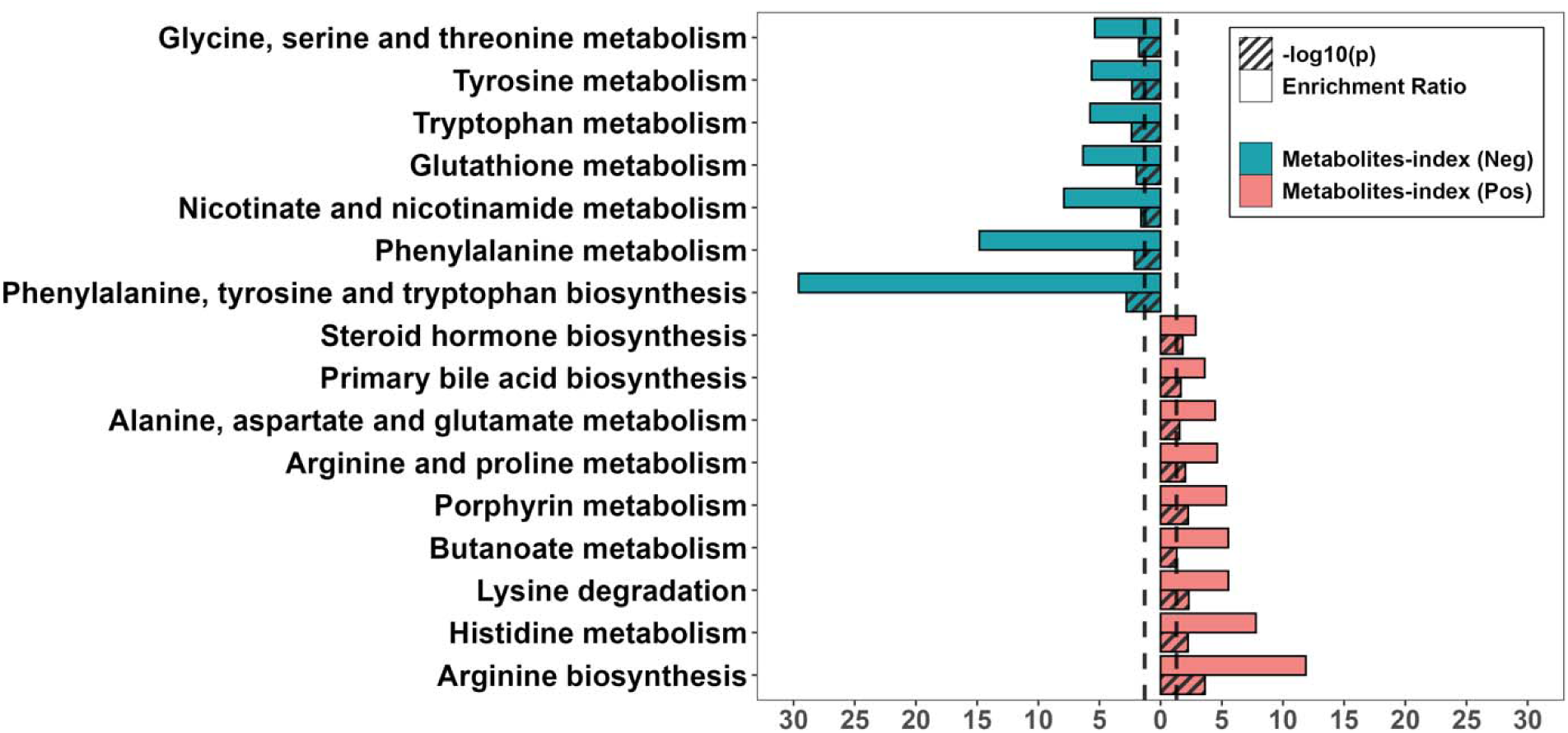
Summary of significantly enriched metabolic pathways (Fisher’s p<0.05) from the KEGG database, analyzed using MetaboAnalyst’s overrepresentation analysis (ORA). The turquoise bars represent the pathway analysis results using 57 influential metabolites exceeding the cutoff threshold in the metabolites-index (Neg), while the red bars represent results based on 72 influential metabolites from the positive metabolites-index (Pos). The X-axis displays the identified enriched metabolic pathways, while the Y-axis indicates the corresponding enrichment ratios or -log10 p-values. A higher enrichment ratio suggests a stronger overrepresentation of that pathway.

To determine whether the mediation effects observed in the PFAS-index were consistent with individual PFAS, we extended our analysis to investigate the mediation effects of individual PFAS, focusing specifically on PFHpS and PFHxS, as they contributed most prominently to the calculated PFAS-index (see **Figure S4)** and showed a significant association with IgG levels (see **Figure S5)**. The metabolites-index (Neg) demonstrated a significant mediation effect on the association between PFHxS and IgG levels (see **Figure S8**). The total effect was -0.741 (95% CI: -1.263, -0.217), with the metabolites-index (Neg) showing a significant indirect effect of - 0.632 (95% CI: -1.237, -0.065) and the metabolites-index (Pos) showing a borderline significant indirect effect of -0.518 (95% CI: -1.035, 0.025) in the analysis of single mediator. In the joint mediation analysis, the metabolites-index (Neg) remained significant with an effect of -0.299 (p = 0.047) while the metabolites-index (Pos) remained a borderline significant effect of -0.360 (p = 0.069). Similar findings were observed for PFHpS (**Figure S9**), where the total effect was -0.937 (95% CI: -1.683, -0.287), the metabolites-index (Neg) showing a significant indirect effect of - 0.671 (95% CI: -1.338, -0.097) while the metabolites-index (Pos) showing a borderline significant indirect effect of -0.597 (95% CI: -1.269, 0.014) in the analysis of single mediator. In the joint mediation analysis, the metabolites-index (Neg) remained significant with an effect of - 0.315 (p = 0.046), while the metabolites-index (Pos) had a borderline significant effect of -0.416 (p = 0.082).

## DISCUSSION

We demonstrate a novel application of WQS in mediation analysis to unravel biological pathways and important metabolites that underlie the previously reported inverse relationship between PFAS exposure and SARS-Cov-2 IgG titer levels. Our results showed that exposure to a PFAS mixture may be associated with alterations in the balance of metabolites, ultimately contributing to a reduced plasma IgG production. This involved two potential mechanisms: increasing metabolite levels linked to lower IgG levels and decreasing metabolite levels associated with higher IgG levels, through distinct and possibly overlapping metabolic pathways. The aim of this approach was to reduce bias due to collinearity and estimate the cumulative effect of plasma metabolites associated with IgG levels in both positive and negative directions. As a result, by constructing metabolites indices, we reduced the dimensionality of 286 metabolites to 2, thereby lowering the computational costs and avoiding multiple comparison errors.

The significant mediation effects of the positive and negative metabolites-indices in both single and multiple mediators analysis suggests that the exposure influences the outcome through different sets of metabolites. Some metabolites may worsen the outcome while others may have a protective effect. In particular, the PFAS-index increases a set of metabolites that are associated with lower SARS-CoV-2 IgG titer levels suggesting that the PFAS-index could be leading to a reduction in IgG levels through this set of metabolites [metabolites index (Neg)]. The PFAS-index also decreases a set of metabolites that are associated with higher IgG titer levels suggesting that the exposure could be suppressing metabolites that would normally boost the IgG levels [metabolites index (Pos)]. The findings that both metabolites-indices significantly mediate the relationship between PFAS-index and SARS-CoV-2 antibody levels provides insight into how PFAS mixture exposure may influence immune function through multiple metabolic pathways. Specifically, the two indices represent distinct sets of metabolites with different patterns: one index captures metabolites whose levels decrease in association with IgG, while the other captures those that increase. This suggests that PFAS exposure may simultaneously activate or suppress different metabolic processes, each impacting the immune response in a unique way, and these distinct pathways likely reflect complex biological mechanisms by which PFAS interferes with immune function.

In the analysis of the single mediator, the indirect effects of metabolites-indices (Neg) and (Pos) were analyzed separately. Both indirect effects were found to be significant, indicating that each mediator independently accounts for a portion of the relationship between the PFAS-index and IgG levels. Notably, the significant direct effect observed with the metabolites-index (Pos) suggests that, even after accounting for the mediator’s effect, part of the total effect remains unexplained. This finding implies that PFAS exposure might influence IgG levels through multiple metabolic pathways, each with distinct roles in IgG regulation. Conversely, in the joint mediation model, while the indirect effects for both mediators remained significant, the direct effect became non-significant. This shift suggests that when the effects of both indices are considered together, they collectively explain most of the association between the PFAS-index and IgG levels. Specifically, the single mediator analysis showed that 74.6% (95% CI: 45.9%, 98.0%) of the effect of PFAS-index on IgG levels was mediated by the metabolites-index (Neg), and 68.6% (95% CI: 41.8%, 94.1%]) was mediated by the metabolites-index (Pos). However, in the joint mediation analysis, adjusting for both metabolites-indices simultaneously revealed a cumulative mediation effect of 83.8% (95% CI: 58.1%, 98.7%]). These results suggest that there is likely overlap between the pathways through which each index mediates the link between PFAS-index and IgG levels (VanderWeele TJ, 2014).

The pathway analysis revealed that the majority of significantly enriched pathways were related to amino acid metabolism. The most significantly enriched pathway in the metabolites- index (Neg) was phenylalanine, tyrosine, and tryptophan biosynthesis. We found that the two metabolites involved in this pathway, phenylalanine and tyrosine, both had relatively high weights in the index (ranked 8 and 3), and both are associated with COVID-19 (Troisi J, et al., 2020). Troisi et al. demonstrated an increasing trend in IgG levels from controls to asymptomatic and mild severity patients, whereas tyrosine and phenylalanine concentrations were lower in mildly symptomatic patients compared to both controls and asymptomatic individuals. Additionally, phenylalanine, tyrosine, and tryptophan biosynthesis has been reported as one of the most common metabolic pathways in metabolomics studies of PFAS exposure (Guo P, 2022, et al., 2024).

Our results also suggest that tryptophan metabolism plays a biological role linking PFAS exposure to IgG titer levels. Metabolites involved in tryptophan metabolism, such as tryptamine, tryptophan, indole-3-pyruvate, and indole-3-acetic acid, were moderately weighted in the metabolites-index (Neg) (ranked 11, 22, 31, and 52), indicating that elevated levels of these metabolites are associated with decreased IgG levels. These findings align with previous studies (Thomas T, et al., 2020; Gardinassi LG,et al., 2020; Takeshita H, et al., 2022; Valdés A, et al.,2022) where altered tryptophan metabolism is considered one of the top pathways affected in COVID-19 cases. Specifically, decreased serum tryptophan and indole-3-pyruvate levels were found in COVID-19 patients compared to healthy individuals (Thomas T, et al., 2020), and tryptophan levels were also found to be decreased in post-acute sequelae of COVID-19 (PASC, “Long COVID”) patients (Wong AC, et al., 2023). Furthermore, tryptophan metabolism is one of the most reported PFAS-associated metabolic pathways (Guo P, et al., 2022).

Arginine metabolism plays an important role in immune response and was the most significantly enriched pathway in the metabolites-index (Pos). Arginine was highly weighted (ranked 9) in the metabolites-index (Pos). The levels of arginine available in the body significantly influence the normal functioning of the immune system (Adebayo A, et al., 2021), particularly via supporting the proliferation of T cells (Zhu X, et al., 2014). Exposure to PFAS mixtures may decrease the plasma arginine levels, potentially leading to reduced T cell proliferation and diminished T cell memory response, which results in a weaker immune response, including lower IgG levels (Congcong Liu, 2023).

Finally, while the significant pathways contributing to the two metabolites indices were distinct, the indirect effects from the joint mediator analysis decreased after accounting for the other index. This suggests that there is some contribution from overlapping pathways, even if those pathways did not reach significance in the Fisher’s exact test. For example, taurine and hypotaurine metabolism were shared between the two indices but not significantly enriched in both, whereas pathways such as tryptophan metabolism and arginine biosynthesis were significantly enriched in one index but not the other. For instance, primary bile acid biosynthesis was shared between the two indices but only significantly enriched in the metabolites-index (Pos). The presence of primary bile acid-related metabolites in the metabolites-index (Pos) (such as cholesterol, chenodeoxycholic acid, taurine, and cholic acid) and in the metabolites-index (Neg) (including glycine and taurocholate) suggests that the primary bile acids pathway has a complex, dual role in relation to IgG levels. Bile acids are known to impact gut microbiome health and microbial balance, thereby influencing key pathways involved in immune regulation and inflammation (Fiorucci S,et al., 2018). Clinical studies have revealed alterations in bile acid metabolism in COVID-19 patients (Valdés A,et al., 2022). Certain bile acids, such as chenodeoxycholic acid (CDCA), have been shown to activate nuclear receptors like the Farnesoid X receptor (FXR) and the G-protein coupled receptor (TGR5), which regulate glucose, lipid, and energy metabolism and are involved in inflammation and immune responses (Chiang JYL,et al., 2020). However, CDCA has also been found to inhibit IgG production while enhancing intracellular IgG concentration (Correia L, et al., 2001). Furthermore, exposure to PFAS may be associated with alterations in the balance of bile acids, altering the metabolic profile in ways that could impair immune function (Guo P, et al., 2022). Individual PFAS increases cholesterol levels and enhances bile acid uptake in the gut, potentially promoting inflammation and leading to cellular stress and damage that could impair immune responses (Behr AC, et al., 2020; Salihović S, et al., 2019). This may partially explain the overlapping mediation effects observed.

In the mediation analysis examining the relationship between metabolite indices, individual PFAS compounds, and IgG levels, we found a significant mediation effect of the metabolites- index (Neg) for both PFHxS and PFHpS on IgG levels (see **Figure S8** and **Figure S9**). In contrast, the metabolites-index (Pos) showed only borderline significant indirect effects (p = 0.054 and p = 0.069). This may be due to the limited sample size or may suggest that individual PFAS compounds lack the statistical power to detect positive mediation effects. This supports our rationale for using overall mixture effects in mediation analysis, reinforcing the idea that exposome influences on metabolites are subtle and require large datasets to detect small but meaningful associations. WQS regression aids this process by summarizing PFAS exposures into a single index that captures the most relevant variation even when individual associations are weak. Further research with larger sample sizes are needed to confirm the mediation effects of the metabolites-indeces for individual PFAS on IgG levels.

The primary limitation of this study is its small sample size, which reduces statistical power, increases the risk of inflated effect sizes, and limits the ability to examine complex relationships or perform subgroup analyses. These constraints should be considered when interpreting the findings. To address high collinearity in the high-dimensional metabolomics data, we applied WQSrs, an ensemble random subset approach that provides robust parameter estimation in complex mixtures. This method ensures stable performance across varying conditions of collinearity and predictor set size. While WQS has been criticized for potential bias (Keil AP, et al., 2020), 2i-WQS (Renzetti S, et al., 2023), an extension of WQS, enables the simultaneous estimation of both positive and negative mixture effects within a single model, similar to Quantile G computation (Keil AP, et al., 2020). However, without a random-subset implememntation, its ability to handle complex correlation structures in high-dimensional datasets remains untested. Future research integrating 2i-WQS with a random subset implementation could significantly advance high-dimensional mixture analyses, especially when the mixture has bidirectional effects on the outcome. Bayesian Kernel Machine Regression (BKMR, Bobb JF, et al., 2014) was also considered, as it is well-suited for modeling nonlinear and interaction effects among exposures. It has been incorporated into a causal mediation framework (Devick KL, et al., 2022) to account for nonlinear and interactions effects among exposures, confounders, mediator, and outcome. However, its application to high-dimensional mediators has yet to be tested, and the complexitiy of results makes interpretation challenging. Even though PFHxS and PFHpS have biological half-lives of 1.5 and 2.9 years, respectively (Xu Y, et al., 2020; Nicole W., 2020), our study is cross-sectional by nature and no causal inference can be made (Winer ES, et al., 2016). Nonetheless, as a preliminary investigation, the key metabolic pathways identified in our two metabolite indices have been thoroughly studied in previous research on PFAS exposures and SARS-CoV-2, and our findings are consistent with those studies. Further studies should consider increasing the sample size and including more diverse cohorts to enhance the statistical power and improve the robustness of results.

## CONCLUSIONS

In summary, our study presented a novel application of WQS in mediation analysis, simplifying the complexity of high-dimensional metabolomics data by consolidating them into two indices. Our analysis demonstrates that PFAS exposures may reduce IgG levels through potential mechanisms that involve both increasing metabolite levels linked to lower IgG levels and decreasing metabolite levels associated with higher IgG levels, with distinct and overlapping pathways contributing to these effects. To confirm these results and gain a deeper understanding of the causal mechanisms, future research should include larger, more diverse cohorts with longitudinal designs. Overall, these results suggest potential biological pathways in support of immunotoxicity of PFAS. This study offers a valuable analytical approach for identifying the metabolic effects linking exposure-outcome associations to enhance causal inference in observational studies.

## CRediT authorship contribution statement

Haibin Guan: Conceptualization, Methodolohgy, Formal analysis, Visualization, Writing- original draft, Writing- review & editing. Jia Chen: Conceptualization, Funding acquisition, Methodology, Formal analysis, Supervision, Writing- original draft, Writing- review & editing. Kirtan Kaur: Writing- review & editing. Bushra Amreen: Writing- review & editing. Corina Lesseur: Writing- review & editing. Georgia Dolios: Investigation, Writing- review & editing. Syam S. Andra: Writing- review & editing. Srinivasan Narasimhan: Writing- review & editing. Divya Pulivarthi: Writing- review & editing. Vishal Midya: Writing- review & editing. Lotje D. De Witte: Writing- review & editing. Veerle Bergink: Writing- review & editing. Anna-Sophie Rommel: Writing- review & editing. Lauren M. Petrick: Conceptualization, Funding acquisition, Methodology, Formal analysis, Supervision, Writing- original draft, Writing- review & editing.

## Acknowledgments

The authors thank the Generation C team, for collecting and processing biospecimens; Dr. Florian Krammer and his laboratory members for the COVID-19 antibody assay and results.

## Funding acknowledgements and disclaimer

We are grateful for support from the National Institutes of Health grants P30ES023515 and UL1TR004419 and HHEAR U2CES030859 and U2CES026561 and R01HD109613. The Generation C study was partially funded by the US Centers for Disease Control and Prevention (CDC) (contract 75D30120C08186). The NIEHS Human Health Exposure Analysis Resource (HHEAR) NIH 2U2CES026561 funded the PFAS measurements, which were carried out within the Senator Frank R. Lautenberg Environmental Health Sciences Laboratory at the Icahn School of Medicine at Mount Sinai. The NIEHS T32HD049311 funded by K. Kaur.

## Appendices. Supplementary data

Overlap of detected metabolites among RPN, RPP and ZHP data modes (**Figure S1**); Flowchart showcases the mediation analysis workflow conducted in this study (**Figure S2**); Heatmap of pairwise Pearson correlation coefficients among 286 maternal plasma metabolites (**Figure S3**); Summary statistics of 286 metabolites on log2 transformed abundance grouped by the metabolite’s superclass (**Table S1**); Summary statistics for 286 metabolites, including mean, standard deviation, median, and range of log2-transformed abundance, along with their chemical superclass, class, m/z, RT, formula, and detection mode (**Table S2**); Descriptive statistics for the WQS indices and immunoglobulin G (IgG) levels across the study population (**Table S3**); The WQS regression model revealed associations of mixed annotated metabolites with IgG levels (**Table S4**); Associations between PFAS-index and SARS-CoV-2 IgG antibody level (**Figure S4**); Forest plot of the univariate linear regression analysis for individual PFAS (**Figure S5**); Bar graph shows the weights of metabolites that surpassed the cutoff threshold for the metabolites- indices negatively and positively associated with IgG levels (**Figure S6**); Summary of all identified metabolic pathways from the KEGG database, analyzed using MetaboAnalyst’s overrepresentation analysis (ORA) (**Figure S7**). Mediation analysis of metabolites-indices on PFHxS and IgG levels (**Figure S8**); Mediation analysis of metabolites-indices on PFHpS and IgG levels (**Figure S9**);

## Data Availability Statement

The maternal plasma PFAS measurements and SARS-CoV-2 IgG antibody quantification data have been published previously (Kaur et al., 2023). The maternal plasma metabolite dataset, derived from LC-HRMS analysis, was specifically generated for this study. All datasets produced and analyzed during the current study are available upon request from the corresponding author.

