## Supplementary material for "High-dimensional mediation analysis to elucidate the role of metabolites in the association between PFAS exposure and reduced SARS-CoV-2 IgG in pregnancy": SI: SI.docx

**Figure S1**. Overlap of detected metabolites among RPN, RPP and ZHP data modes

**Figure S2**. Flowchart showcases the mediation analysis workflow conducted in this study

**Figure S3**. Heatmap of pairwise Pearson correlation coefficients among 286 maternal plasma metabolites

**Table S1**. Summary statistics of 286 metabolites on log2 transformed abundance grouped by the metabolite’s superclass

**Table S2**. Summary statistics for 286 metabolites, including mean, standard deviation, median, and range of log2-transformed abundance, along with their chemical superclass, class, m/z, RT, formula, and detection mode

**Table S3.** Descriptive statistics for the Weighted Quantile Sum (WQS) indices and immunoglobulin G (IgG) levels across the study population (N=59)

**Table S4**. The WQS regression model revealed associations of mixed annotated metabolites with IgG levels

**Figure S4**. Associations between PFAS-index and SARS-CoV-2 IgG antibody level

**Figure S5.** Forest plot of the univariate linear regression analysis for individual PFASs

**Figure S6**. Bar graph shows the weights of metabolites that surpassed the cutoff threshold for the metabolites-indices negatively and positively associated with IgG levels

**Figure S7**. Summary of all identified metabolic pathways from the KEGG database, analyzed using MetaboAnalyst's overrepresentation analysis (ORA)

**Figure S8.** Mediation analysis of metabolites-indices on **PFHxS** and IgG levels

**Figure S9.** Mediation analysis of metabolites-indices on **PFHpS** and IgG levels

**Figure S1**. Overlap of detected metabolites among RPN, RPP and ZHP data modes. A) The venn diagram illustrates the intersection of metabolites detected in three different data modes: RPN, RPP and ZHP, prior to filtering. B) The venn diagram illustrates the intersection of metabolites detected in three different data modes: RPN, RPP and ZHP^1^, after filtering. C) Following filtering, the distribution of 286 measured metabolites categorized by chemical superclass searched by using “ClassyFire”.

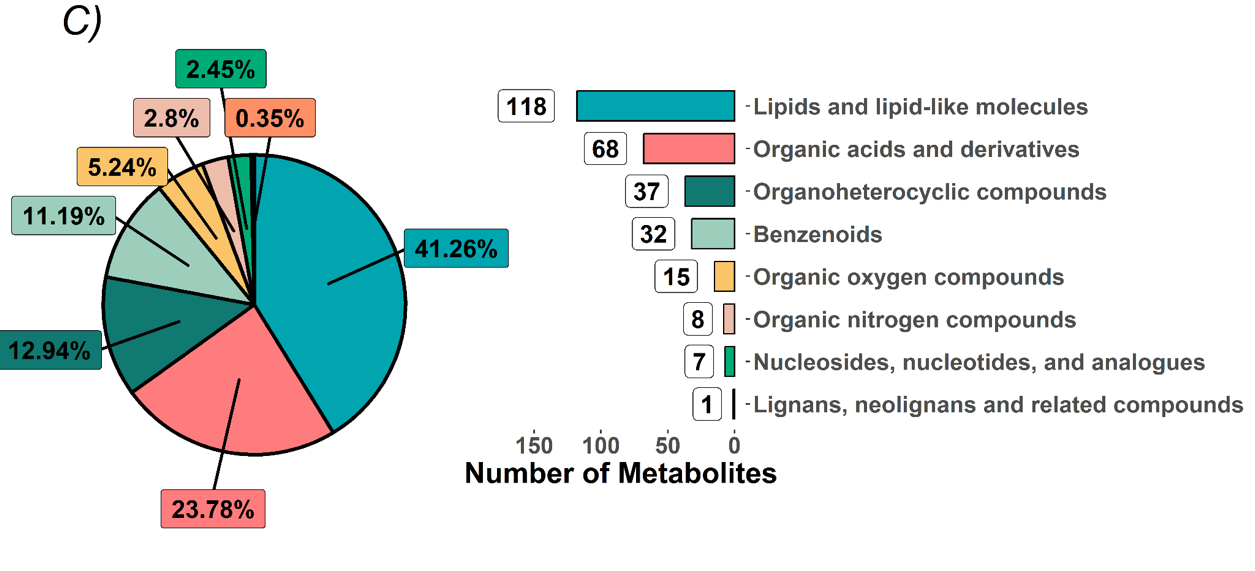

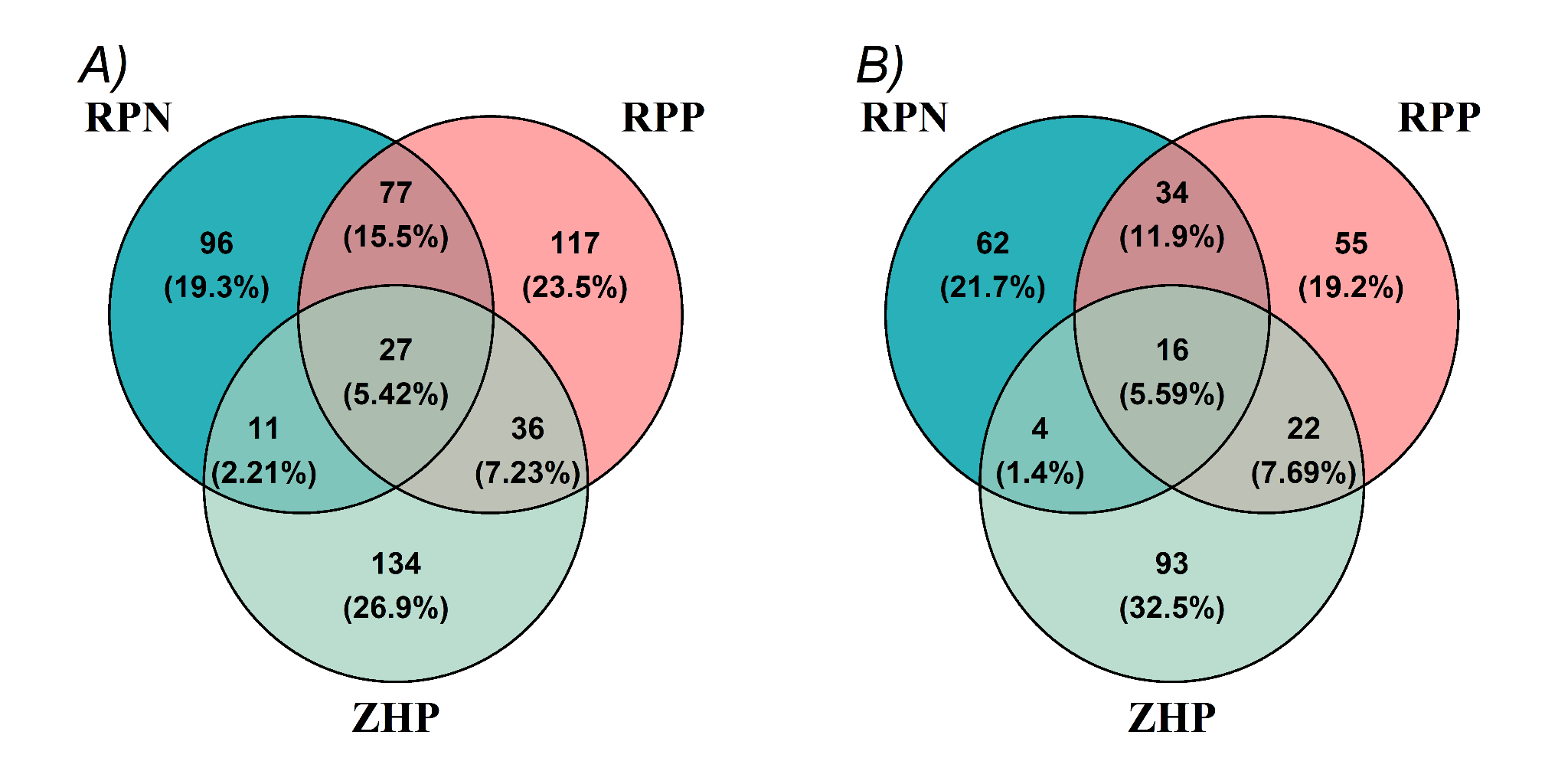

**^1^RPP:** Reverse Phase Liquid Chromatography (RPLC) in Positive ionization mode; **RPN:** Reverse Phase Liquid Chromatography (RPLC) in Negative ionization mode; **ZHP:** Hydrophilic Interaction Liquid Chromatography (HILIC) in Positive ionization mode.

**Figure S2**. Flowchart showcases the mediation analysis workflow conducted in this study.

**Top:** Single mediator analysis with metabolites-index (Neg) or metabolites-index (Pos).

Total effect of PFAS-index on IgG levels, without taking mediator into account - path $c$

PFAS-index is hypothesized to influence IgG levels through metabolites-index(Neg) or metabolites-index(Pos) independently - path $\alpha_{1}\beta_{1}$ or $\alpha_{2}\beta_{2}$; after accounting for the individual mediator, the remaining direct effect on IgG levels - path $c_{1}'$ or $c_{2}'$

**Bottom:** Multiple mediators analysis with both metabolites-index (Neg) and metabolites-index (Pos) included in one model.

Total effect of PFAS-index on IgG levels, without taking mediators into account - path $c$

PFAS-index is hypothesized to influence IgG levels indirectly through two mediators: metabolites-index(Neg) and metabolites-index(Pos), in combination - path $\alpha_{1}{\beta'}_{1}$ or $\alpha_{2}{\beta'}_{2}$; after accounting for the combined influence of both mediators, the remaining direct effect - path $c'$

All models were adjusted for maternal age, maternal race/ethnicity, parity, type of insurance, gestational age at blood draw (weeks), receipt of the COVID-19 vaccine before blood collection, and pre-pregnancy BMI.

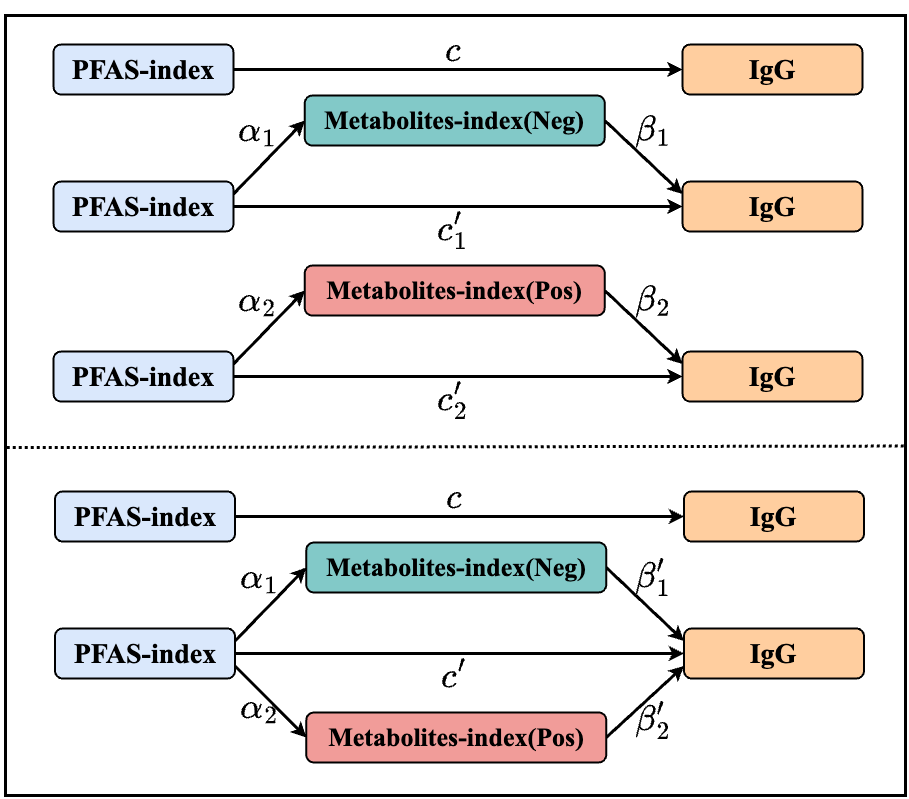

WQS regression model

$IgG= c_{0}+c\times\sum_{j=1}^{m} w_{j}p_{j}+\sum_{i=1}^{k} c_{i}Z_{i} +\epsilon_{Y1}$	(1)

We first implemented WQS to identify the total effect of the PFAS-index on the SARS-CoV-2 IgG antibody levels by using the model in equation (1), where $c_{0}$ is the intercept, $c_{i}$ are coefficients for covariate $Z_{i}$ , $\epsilon_{Y}$ is the error term, and the $PFAS.index=\sum_{j=1}^{m} w_{j}p_{j}$ which represents the WQS index that weights and sums the components included in the 9 PFAS exposure mixture. The weight $w_{j}$ indicates the relative importance of each PFAS exposure in the association between PFAS-index and IgG. The estimated $c$ is the parameter summarizing the total effect of the PFAS-index on IgG levels.

Single mediator model

We then implemented random-subset WQS to identify the metabolites-index in the negative and positive directions that describes the influence of the PFAS-index on the metabolite profiles.

$Metabolite-index\left( Neg \right)=\eta_{0}+ \alpha_{1}\times PFAS-\mathrm{index}+\sum_{i=1}^{k} \eta_{i}Z_{i} +\epsilon_{M1}$	(2)

$Metabolite-index\left( Pos \right)={\eta^{'}}_{0}+ \alpha_{2}\times PFAS-\mathrm{index}+\sum_{i=1}^{k} {\eta'}_{i}Z_{i} +\epsilon_{M2}$	(3)

In this mediator model, where the $\eta_{0}$and${\eta'}_{0}$ are intercepts, $\eta_{i}$ and ${\eta'}_{i}$ are coefficients for covariate $Z_{i}$, $\epsilon_{M1}$ and $\epsilon_{M2}$ are error terms, and $Metabolites-index(Neg)=\sum_{j=1}^{n} \theta_{1j}q_{j}$and $Metabolites-index(Pos)=\sum_{j=1}^{n} \theta_{2j}q_{j}$ represent the WQS index that weights and sums the components included in the high dimensional metabolites set in negative and positive direction, respectively. The weights $\theta_{1j}$ and $\theta_{2j}$ indicate the relative importance of each metabolite in the association between PFAS-index and metabolites-index (Neg) and PFAS-index and metabolites-index (Pos), respectively. As such, $a_{1}$ and $a_{2}$ are the parameters summarising the effects of PFAS-index on the two mediators.

We then determine the direct effect of the PFAS-index on IgG titer levels and the indirect effect of PFAS on IgG titer levels through the mediation of metabolites using the following outcome model, respectively:

$IgG=\gamma+c_{1}^{'}\times PFAS-index+ \beta_{1}\times Metabolites-index(Neg)+ \sum_{i=1}^{k} \gamma_{i}Z_{i}+$ $\epsilon$	(4)

$IgG=\gamma^{'}+c_{2}^{'}\times PFAS-index+ \beta_{2}\times Metabolites-index(Pos)+$ $\sum_{i=1}^{k} {\gamma'}_{i}Z_{i}$+ $\epsilon'$	(5)

Where the $\gamma$and$\gamma'$ are intercepts, $\gamma_{i}$ and ${\gamma'}_{i}$ are coefficients for covariate $Z_{i}$, and $\epsilon$ and $\epsilon'$ are error terms. We estimate the effect of the PFAS-index on IgG through the metabolites-index in negative or positive direction (indirect effect = $\alpha_{1}\times\beta_{1}$ or $\alpha_{2}\times\beta_{2}$) and the effect of the PFAS-index on IgG unexplained by the metabolites-index (direct effect = $c_{1}'$ or $c_{2}'$).

$$Proportion of Indirect Effect=\frac{Indirect Effect}{Total Effect}$$

Multple-meidator mediation model

In the second scenario, we conducted a mediation model in parallel design with two mediators, metabolites-index (Pos) and metabolites-index (Neg), on PFAS-index and SARS-CoV-2 IgG antibody levels (N = 59) adjusted for maternal age, maternal race/ethnicity, parity, type of insurance, gestational age at blood draw (weeks), receipt of the COVID-19 vaccine before blood collection, and pre-pregnancy BMI. Mediator models utilized the same equations as in (2) and (3), and the outcome model utilized the following equation:

$IgG=\lambda+c^{'}\times PFAS-index+ {\beta'}_{1}\times Metabolites-index(Neg)+{\beta'}_{2}\times Metabolites-index(Pos)+ \sum_{i=1}^{k} \lambda_{i}Z_{i}+e$ (8)

Where the $\lambda$ is the intercept, $\lambda_{i}$ are coefficients for covariate $Z_{i}$, and $e$ is an error term. We can estimate the effect of the PFAS-index on IgG that affects through metabolites-index in a negative direction and metabolites-index in a positive direction (indirect effect of metabolites-index (Neg) = $\alpha_{1}\times\beta_{1}$; indirect effect of metabolites-index (Pos) = $\alpha_{2}\times\beta_{2}$) and the effect of the PFAS-index on IgG unexplained by the metabolites-index (direct effect=$c'$).

**Figure S3**: Heatmap of pairwise Pearson correlation coefficients among 286 maternal plasma metabolites (turquoise indicates positive correlation, red indicates negative correlation). Metabolites are clustered using hierarchical clustering (complete linkage method) based on correlation distances. The metabolites are further color coded according to their superclass. A closer examination of a subset of the heatmap reveals a group of 50 metabolites that exhibit relatively higher correlations with each other.

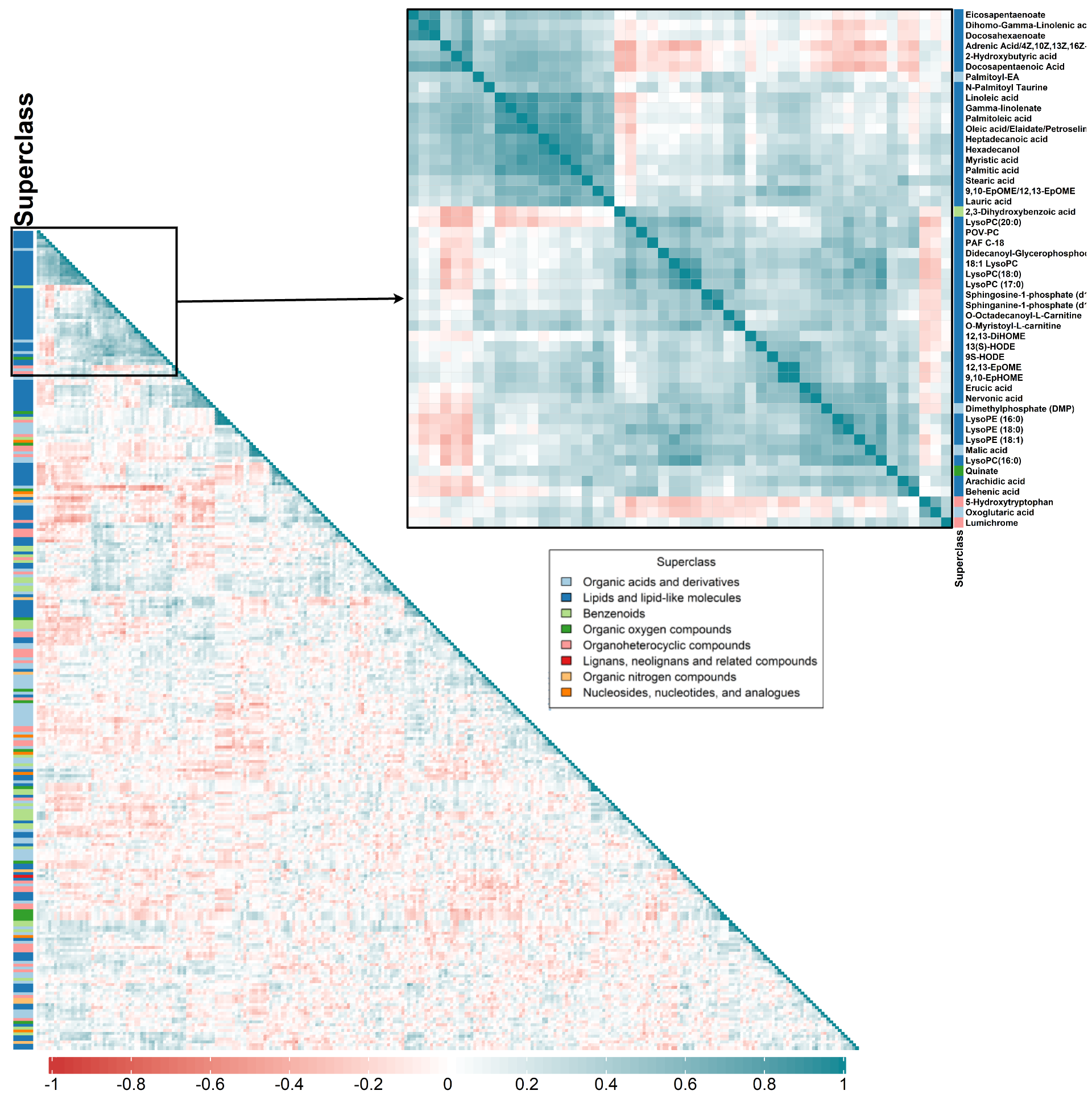

**Table S1**. Summary statistics of 286 metabolites on log2 transformed abundance grouped by the metabolite’s superclass. The median superclass values of the individual metabolite’s medians, 10th percentiles, 90th percentiles, and ratios of 90th to 10th percentiles are reported.

| **Superclass** | **Metabolite** | **Q10 median(min,max)** | **Q90 median(min,max)** | **IQR median(min,max)** | **Min median(min,max)** | **Max median(min,max)** | **Median median(min,max)** | **Mean median(min,max)** | **SD median(min,max)** |
| --- | --- | --- | --- | --- | --- | --- | --- | --- | --- |
| **Benzenoids** | 32 | 14.18 (10.8,21.49) | 15.59 (12,24.06) | 0.61 (0.22,2.2) | 13.75 (10.45,20.68) | 16.43 (12.13,24.73) | 14.88 (11.68,22.84) | 14.83 (11.66,22.88) | 0.48 (0.17,2.05) |
| **Lignans, neolignans and related compounds** | 1 | 18.43 (18.43,18.43) | 19.26 (19.26,19.26) | 0.34 (0.34,0.34) | 18.18 (18.18,18.18) | 19.38 (19.38,19.38) | 18.9 (18.9,18.9) | 18.85 (18.85,18.85) | 0.3 (0.3,0.3) |
| **Lipids and lipid-like molecules** | 118 | 16.3 (9.99,25.09) | 17.9 (12.24,25.8) | 0.79 (0.24,2.33) | 15.73 (9.74,24.94) | 18.56 (12.56,25.89) | 17.33 (11.22,25.47) | 17.27 (11.17,25.43) | 0.61 (0.17,1.63) |
| **Nucleosides, nucleotides, and analogues** | 7 | 13.03 (8.87,18.23) | 13.8 (10.78,19.98) | 0.47 (0.31,0.89) | 12.9 (7.75,17.98) | 14.11 (11.31,20.05) | 13.48 (9.88,19.04) | 13.45 (9.83,19.08) | 0.35 (0.21,0.82) |
| **Organic acids and derivatives** | 68 | 15.43 (11.2,22.28) | 16.23 (12,23.12) | 0.48 (0.19,1.3) | 15.28 (10.86,22.06) | 16.41 (12.18,23.27) | 15.87 (11.6,22.66) | 15.82 (11.62,22.67) | 0.36 (0.15,1.03) |
| **Organic nitrogen compounds** | 8 | 15.53 (12.07,23.67) | 16.31 (13.59,24.34) | 0.49 (0.22,0.79) | 15.4 (11.85,23.5) | 16.41 (14.4,24.47) | 15.92 (12.86,24.01) | 15.92 (12.89,24.01) | 0.33 (0.17,0.61) |
| **Organic oxygen compounds** | 15 | 12.72 (9.53,18.16) | 13.75 (10.47,19.83) | 0.5 (0.21,2.1) | 12.49 (9.13,18.04) | 13.94 (10.74,20.76) | 13.35 (10.07,18.37) | 13.3 (10.02,18.38) | 0.4 (0.16,1.44) |
| **Organoheterocyclic compounds** | 37 | 14.6 (11.35,21.15) | 16.39 (12.25,24.42) | 0.57 (0.22,2.67) | 14.31 (11.15,21.02) | 16.62 (12.35,24.75) | 15.29 (11.81,22.65) | 15.41 (11.81,22.03) | 0.5 (0.17,2.55) |

**Table S2**. Summary statistics for 286 metabolites, including mean, standard deviation, median, and range of log2-transformed abundance, along with their chemical superclass, class, m/z, RT, formula, and detection mode. Additionally, the corresponding weights in two metabolites.indices are provided (SI_TableS2.xlsx file).

**Table S3.** Descriptive statistics for the Weighted Quantile Sum (WQS) indices and immunoglobulin G (IgG) levels across the study population (N=59). The table presents the mean, standard deviation (SD), median, and range (minimum and maximum) for each variable.

|  | **Overall (N=59)** |
| --- | --- |
| **PFAS-index** |  |
| Mean (SD) | 1.45 (0.672) |
| Median [Min, Max] | 1.35 [0.439, 2.56] |
| **Metabolites-index (Neg)** |  |
| Mean (SD) | 1.47 (0.247) |
| Median [Min, Max] | 1.45 [1.04, 2.05] |
| **Metabolites-index (Pos)** |  |
| Mean (SD) | 1.47 (0.263) |
| Median [Min, Max] | 1.45 [0.913, 2.09] |
| **SARS-Cov2 IgG titer** |  |
| Mean (SD) | 2.67 (0.481) |
| Median [Min, Max] | 2.60 [1.70, 3.81] |

**Table S4**. The WQS regression model revealed associations of mixed annotated metabolites with IgG levels.

| **WQS Index** | **Estimate** | **SE** | **P value** |
| --- | --- | --- | --- |
| **PFAS - index** | -0.315 | 0.076 | <0.001 |
| **Metabolites-index (Neg)** | -1.200 | 0.160 | <0.001 |
| **Metabolites-index (Pos)** | 1.249 | 0.140 | <0.001 |

**Figure S4**. **a**) The box plot shows the weights of nine PFAS exposures associated with the PFAS-index. The box-plot depict the 25^th^, 50^th^, and 75^th^ percentiles, while the whiskers extend to the 10^th^ and 90^th^ percentiles. PFAS exposures are ranked based on their median weight, which is indicated as a text label. Diamonds indicate the mean weights, and individual data points represent the estimated weights through the 5000 simulations. All WQS models were adjusted for maternal age, maternal race/ethnicity, parity, type of insurance, gestational age at blood draw (weeks), receipt of the COVID-19 vaccine before blood collection, and pre-pregnancy BMI. The dashed dark line represents the cutoff of 1/9 (equal to the inverse of the number of elements in the mixture as suggested in Carrico et al. 2014). In the current study population (N=59), the largest contributors to this mixture effect in the negative direction are: PFHpS and PFHxS; **b**) A representation of the direction of the association between the PFAS-index vs the SARS-CoV-2 IgG antibody levels (adjusted for the model residual when covariates are included in the model). *** Statistical significance levels for the effects (p < 0.001).

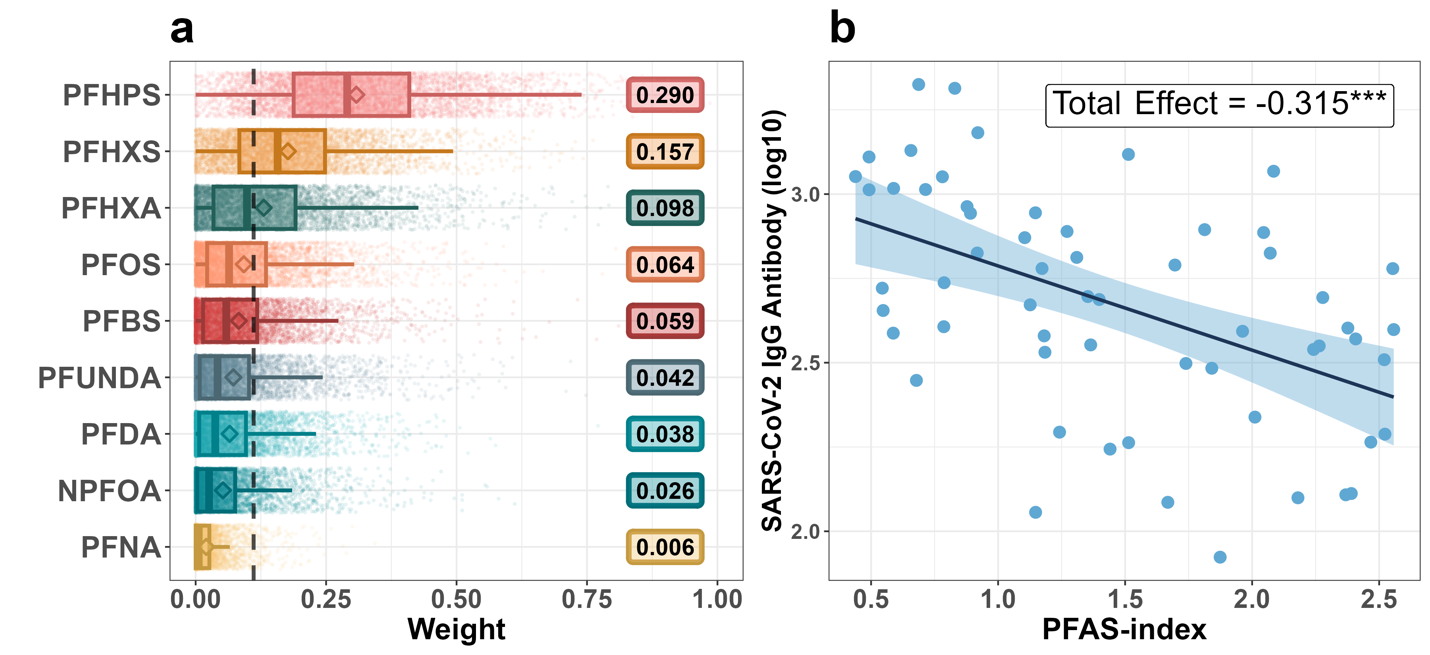

**Figure S5.** Forest plot of the univariate linear regression analysis for individual PFAS, adjusted for maternal age, maternal race/ethnicity, parity, type of insurance, gestational age at blood draw (weeks), receipt of the COVID-19 vaccine before blood collection, and pre-pregnancy BMI.

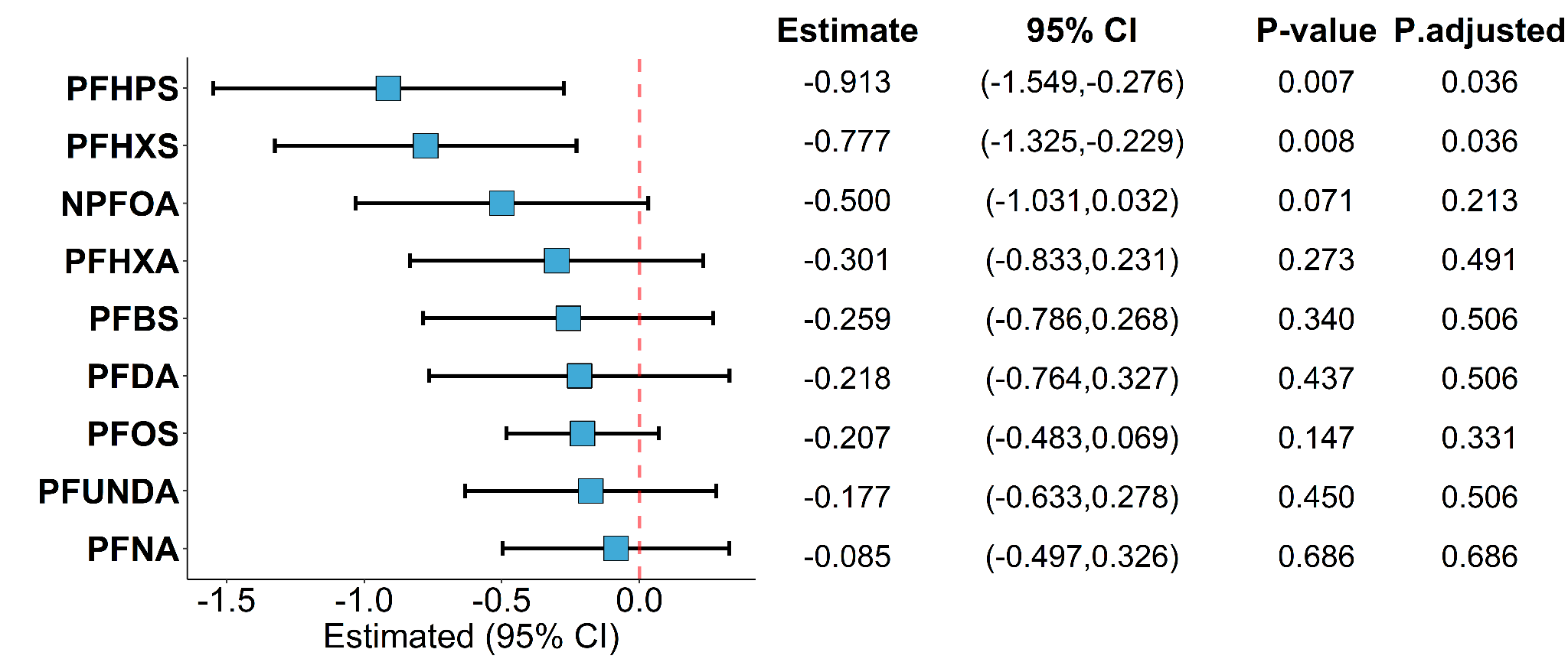

**Figure S6**. The box plot shows the weights of 57 and 72 metabolites whose median weights surpassed the cutoff threshold in metabolites-index (Neg) (**a**) and metabolites-index (Pos) (**b**), respectively. Metabolites are ordered from the highest weight to the lowest. The box-plot depict the 25^th^, 50^th^, and 75^th^ percentiles, while the whiskers extend to the 10^th^ and 90^th^ percentiles. Diamonds indicate the mean weights, and individual data points represent the estimated weights associated with the metabolites-index (Neg) and (Pos) through the 5000 simulations. The dashed dark line represents the cutoff (1/286). Colors represents the respective chemical superclass of each metabolite. Both WQS models were adjusted for maternal age, maternal race/ethnicity, parity, type of insurance, gestational age at blood draw (weeks), receipt of the COVID-19 vaccine before blood collection, and pre-pregnancy BMI.

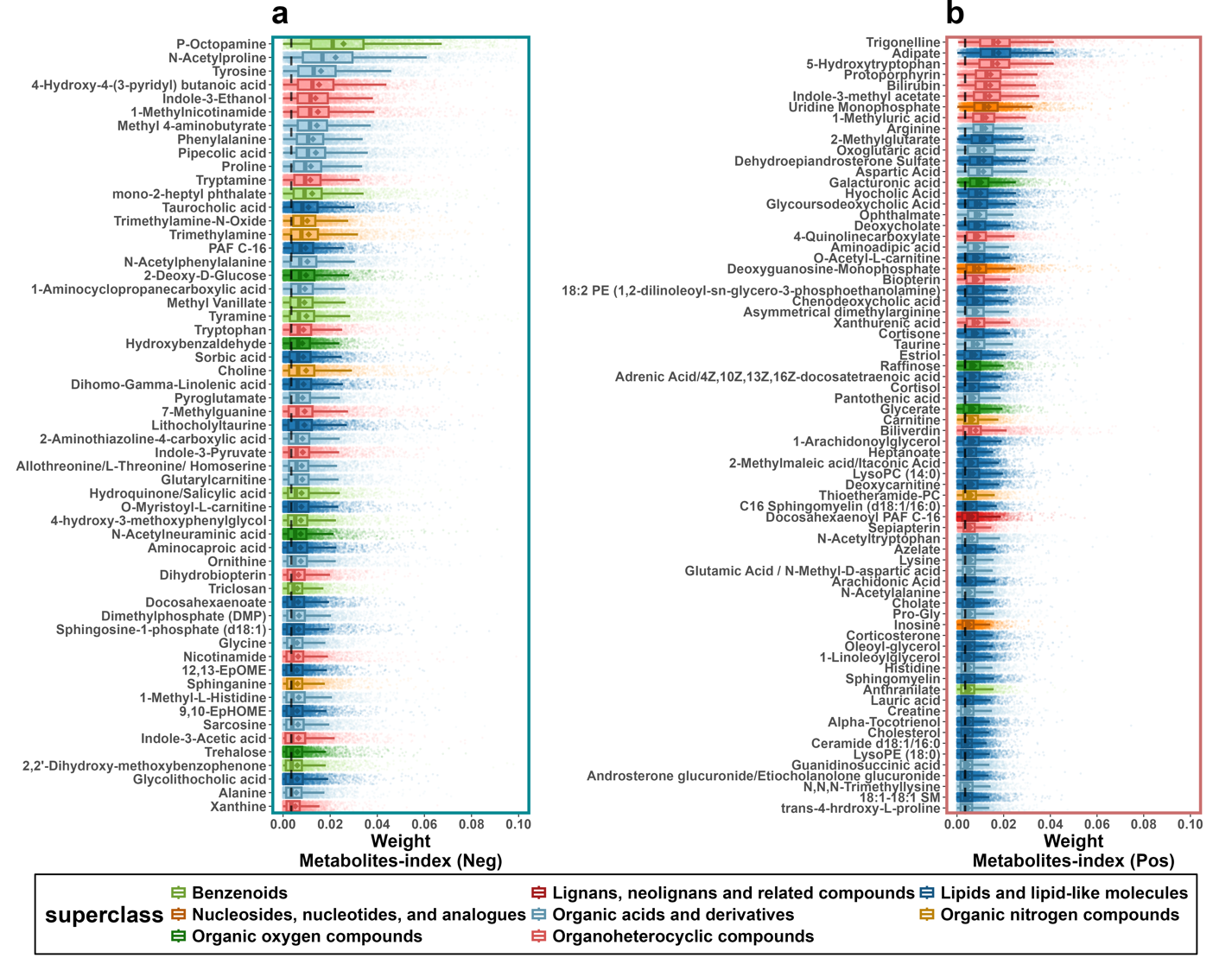

**Figure S7**. Summary of all identified metabolic pathways from the KEGG database, analyzed using MetaboAnalyst's overrepresentation analysis (ORA). The turquoise bars represent the pathway analysis results using 57 influential metabolites exceeding the cutoff threshold in the metabolites-index (Neg), while the red bars represent results based on 72 influential metabolites from the metabolites-index (Pos). The X-axis displays the identified enriched metabolic pathways, while the Y-axis indicates the corresponding enrichment ratios or -log10 p-values^2^.

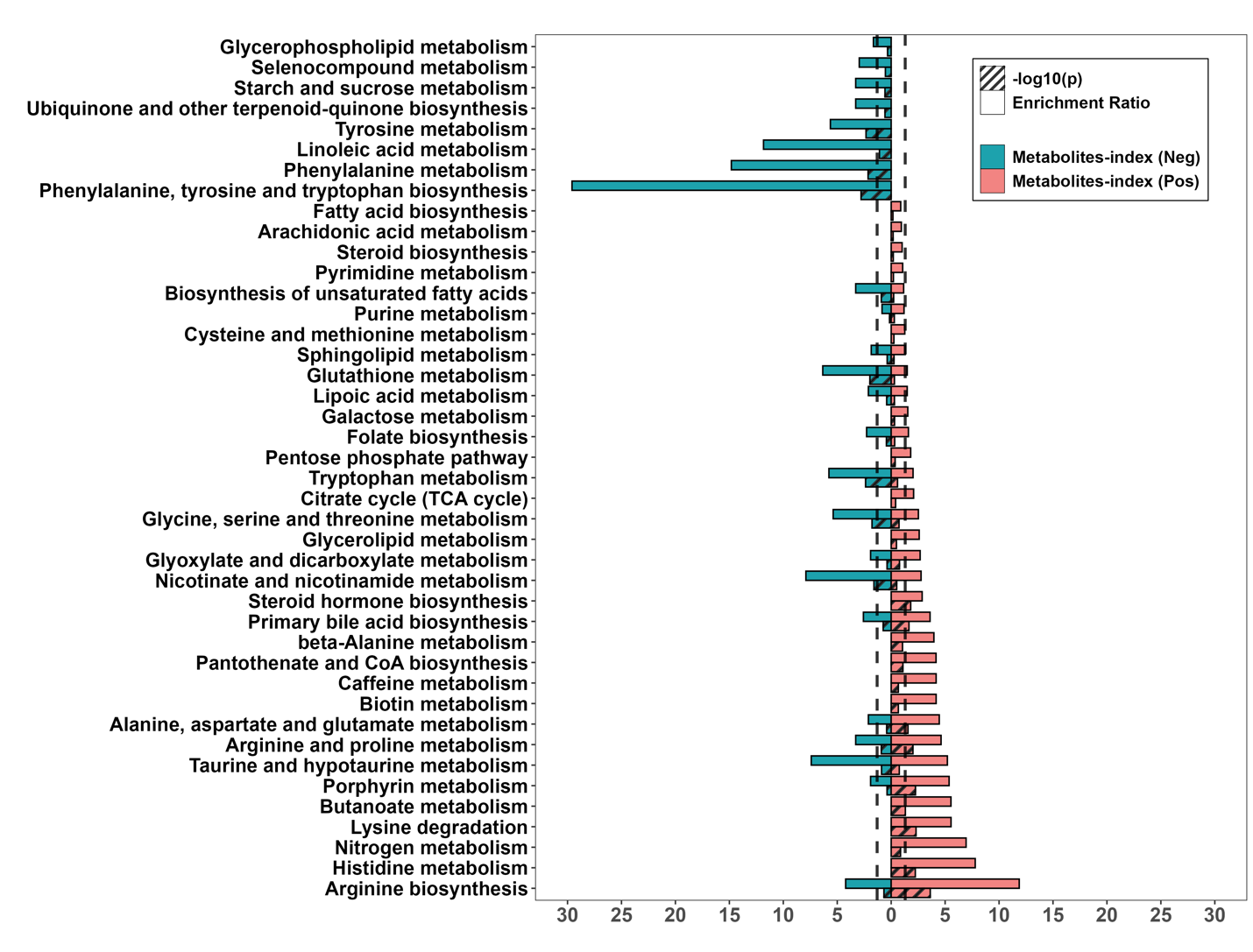

^2^In MetaboAnalyst's Over-Representation Analysis (ORA), the enrichment ratio is calculated as the number of hits within a specific pathway divided by the expected number of hits. A higher enrichment ratio suggests a stronger overrepresentation of that pathway. The corresponding p-value, calculated using Fisher’s exact test, assesses the statistical significance of the enrichment.

**Figure S8.** Analysis results on mediation analysis of metabolites-indices on PFHxS and IgG levels. **a)** Forest plot for mediation analysis of metabolites-index (Neg) on PFHxS and IgG levels (N=59), adjusted for covariates. **b)** Estimated beta from the mediator model for the association between metabolites-index (Neg) and PFHxS, adjusted for covariates. **c)** Estimated beta from the outcome model for the association between IgG levels and metabolites-index (Neg), adjusted for covariates. **d)** Forest plot for mediation analysis of metabolites-index (Pos) on PFHxS and IgG levels (N=59), adjusted for covariates. **e)** Estimated beta from the mediator model for the association between metabolites-index (Pos) and PFHxS, adjusted for covariates. **f)** Estimated beta from the outcome model for the association between IgG levels and metabolites-index (Pos), adjusted for covariates. **g)** Forest plot for joint mediation effect analysis of two metabolites-indices on PFHxS and IgG levels (N=59), adjusted for covariates. All p-values estimated using a two-sided test for t-statistics from linear regression models. Statistical significance levels for the effects: * (p < 0.05) and ***(p < 0.001).

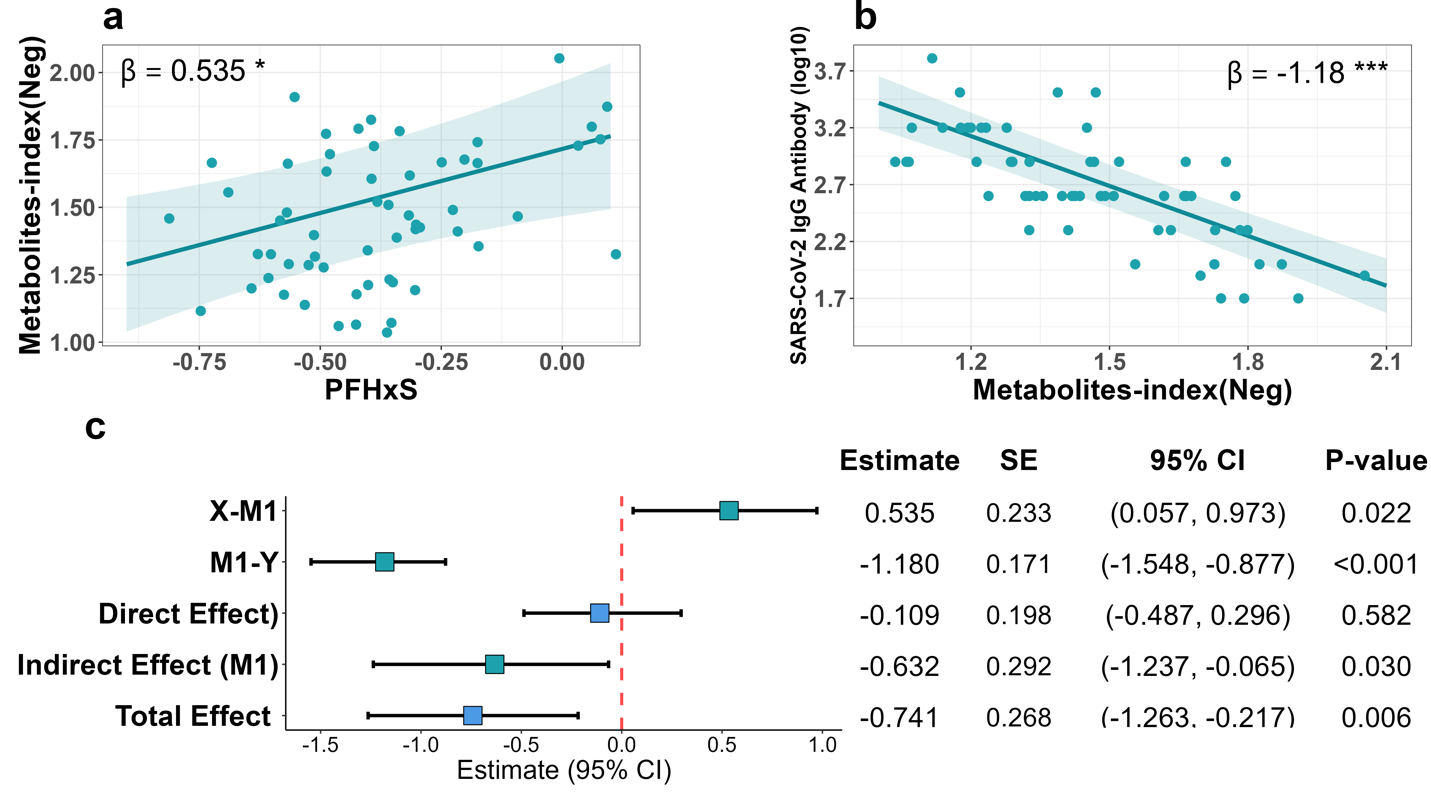

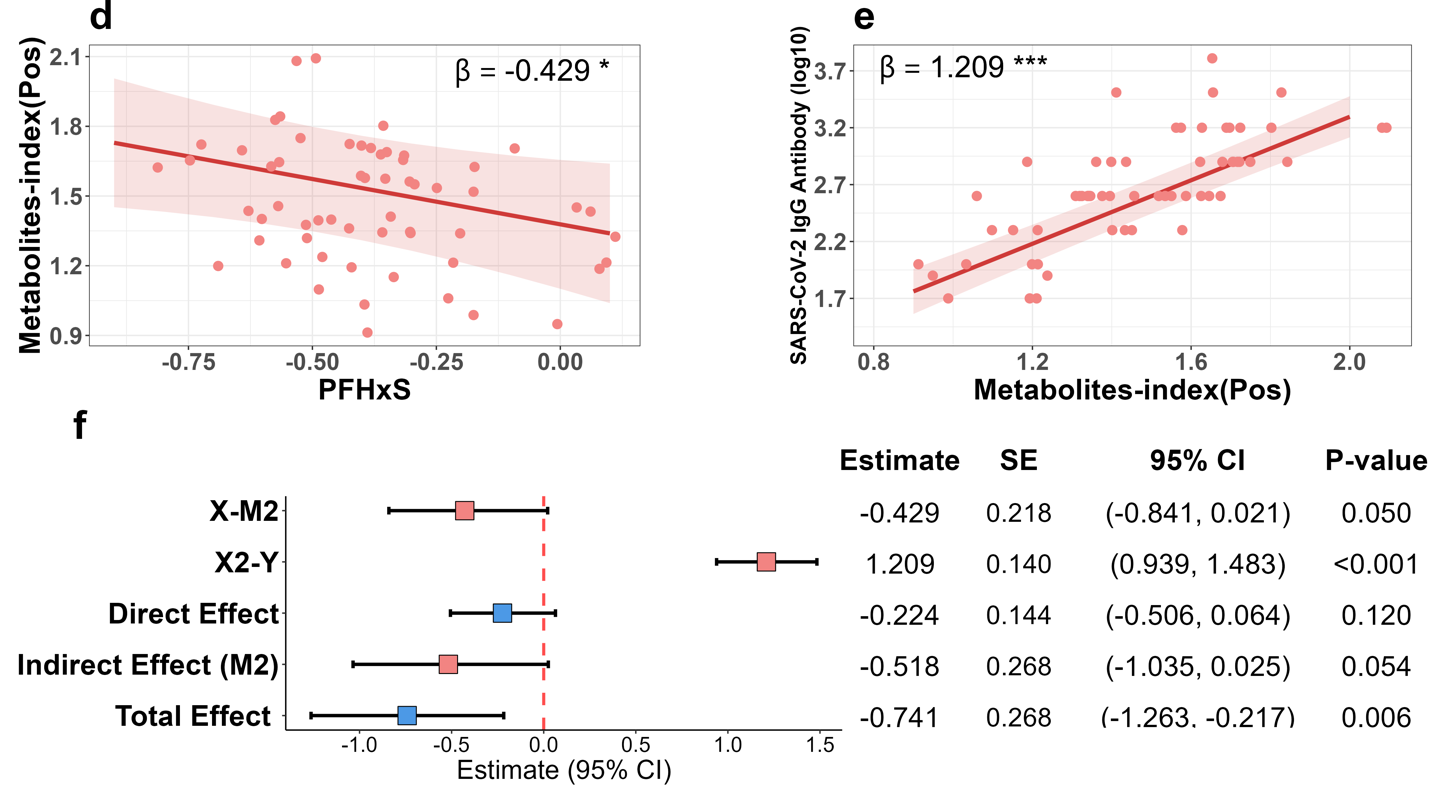

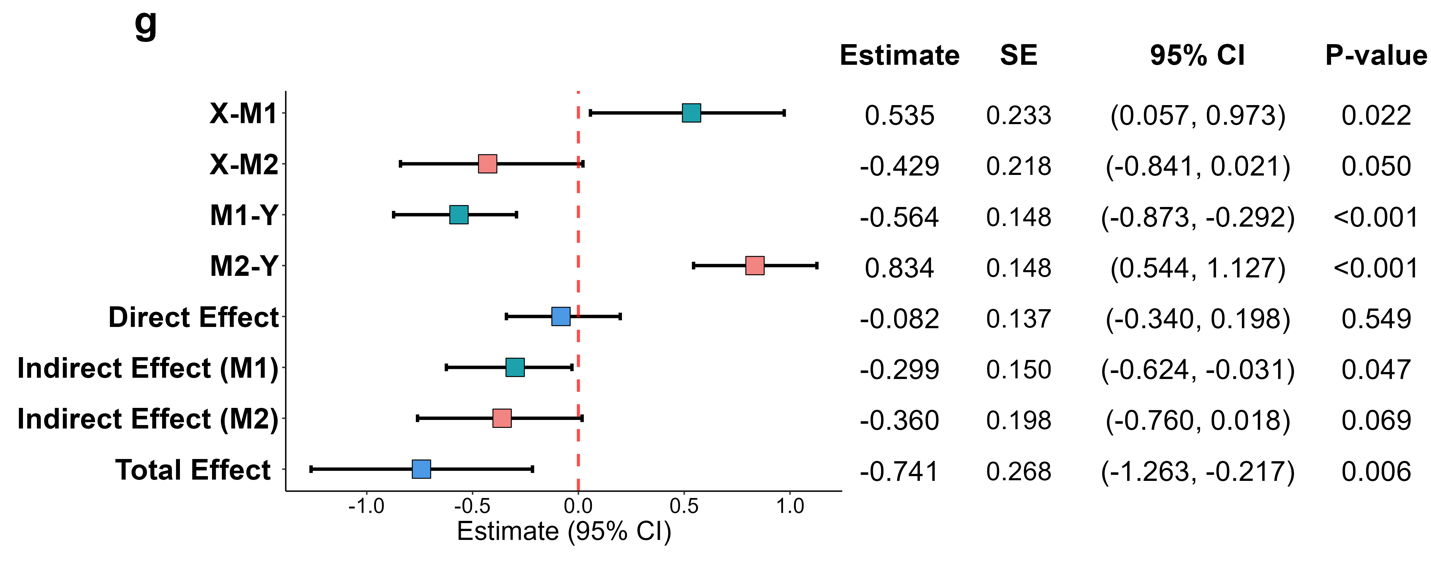

**Figure S9.** Analysis results on mediation analysis of metabolites-indices on PFHpS and IgG levels. **a)** Forest plot for mediation analysis of metabolites-index (Neg) on PFHpS and IgG levels (N=59), adjusted for covariates. **b)** Estimated beta from the mediator model for the association between metabolites-index (Neg) and PFHpS, adjusted for covariates. **c)** Estimated beta from the outcome model for the association between IgG levels and metabolites-index (Neg), adjusted for covariates. **d)** Forest plot for mediation analysis of metabolites-index (Pos) on PFHpS and IgG levels (N=59), adjusted for covariates. **e)** Estimated beta from the mediator model for the association between metabolites-index (Pos) and PFHpS, adjusted for covariates. **f)** Estimated beta from the outcome model for the association between IgG levels and metabolites-index (Pos), adjusted for covariates. **g)** Forest plot for joint mediation effect analysis of two metabolites-indices on PFHpS and IgG levels (N=59), adjusted for covariates. All p-values estimated using a two-sided test for t-statistics from linear regression models. Statistical significance levels for the effects: * (p < 0.05) and ***(p < 0.001)

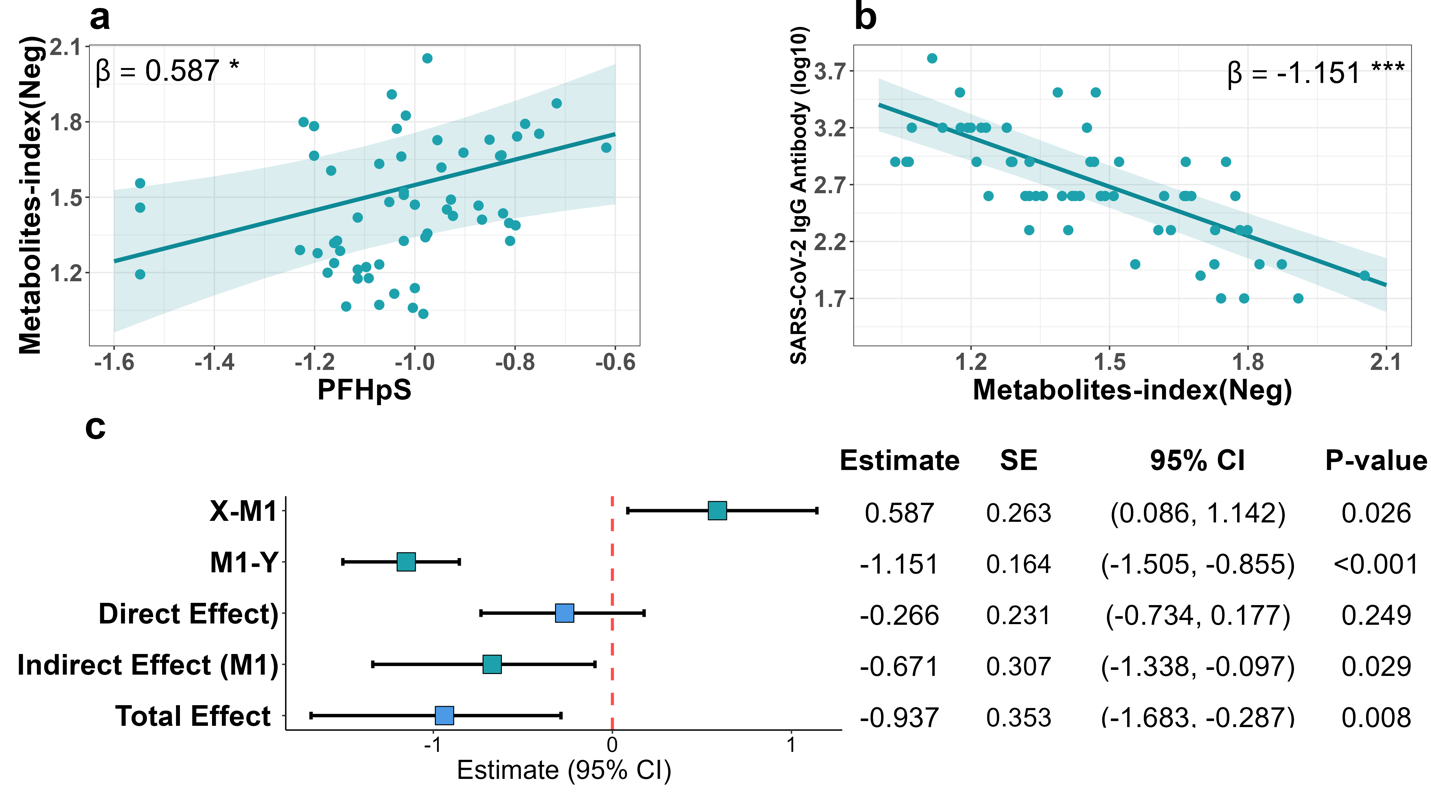

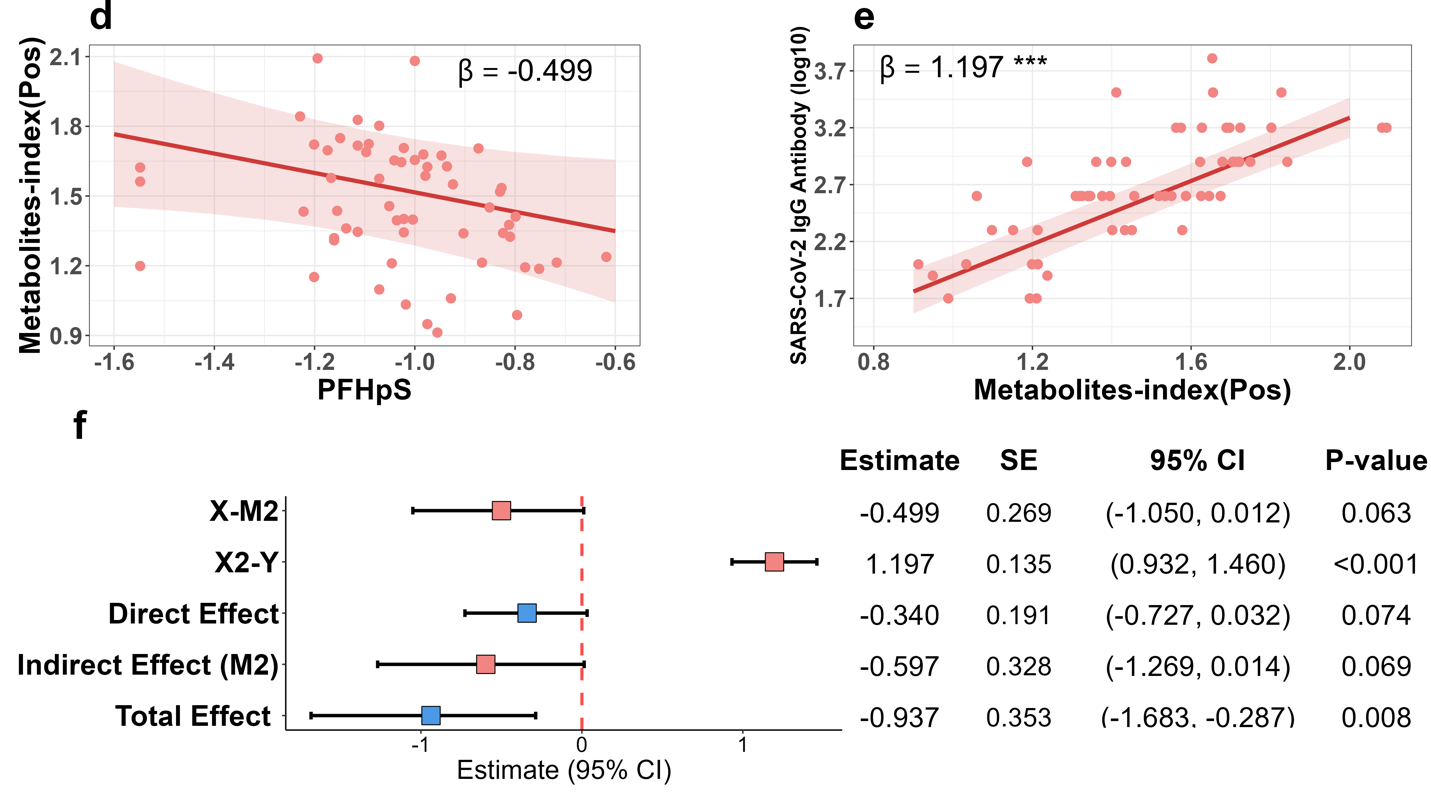

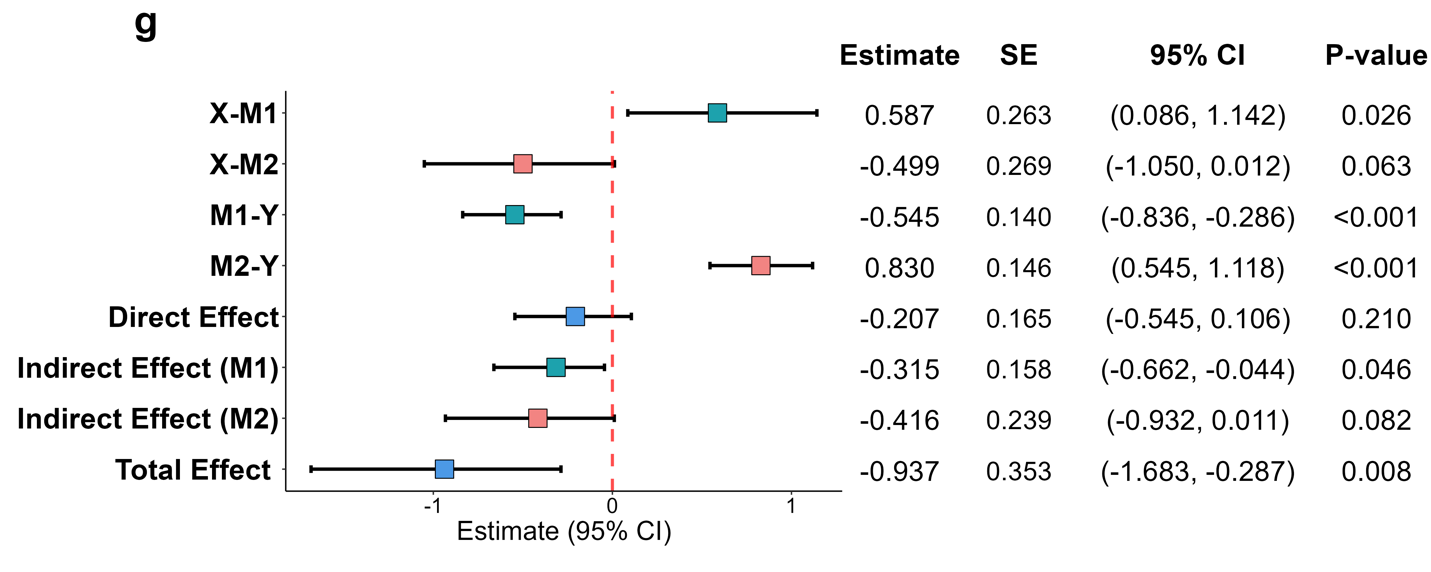
